# A Statistical Pipeline for Identifying Physical Features that Differentiate Classes of 3D Shapes

**DOI:** 10.1101/701391

**Authors:** Bruce Wang, Timothy Sudijono, Henry Kirveslahti, Tingran Gao, Douglas M. Boyer, Sayan Mukherjee, Lorin Crawford

## Abstract

The recent curation of large-scale databases with 3D surface scans of shapes has motivated the development of tools that better detect global patterns in morphological variation. Studies which focus on identifying differences between shapes have been limited to simple pairwise comparisons and rely on pre-specified landmarks (that are often known). We present SINATRA: the first statistical pipeline for analyzing collections of shapes without requiring any correspondences. Our novel algorithm takes in two classes of shapes and highlights the physical features that best describe the variation between them. We use a rigorous simulation framework to assess our approach. Lastly, as a case study, we use SINATRA to analyze mandibular molars from four different suborders of primates and demonstrate its ability recover known morphometric variation across phylogenies.

## Introduction

Sub-image analysis is an important open problem in both medical imaging studies and geometric morphometric applications. The problem asks which physical features of shapes are most important for differentiating between two classes of 3D images or shapes such as computed tomography (CT) scans of bones or magnetic resonance images (MRI) of different tissues. More generally, the sub-image analysis problem can be framed as a regression-based task: given a collection of shapes, find the properties that explain the greatest variation in some response variable (continuous or binary). One example is identifying the structures of glioblastoma tumors that best indicate signs of potential relapse and other clinical outcomes [1]. From a statistical perspective, the sub-image selection problem is directly related to the variable selection problem — given high-dimensional covariates and a univariate outcome, we want to infer which variables are most relevant in explaining or predicting variation in the observed response.

Framing sub-image analysis as a regression presents several challenges. The first challenge centers around representing a 3D object as a (square integrable) covariate or feature vector. The transformation should lose a minimum amount of geometric information and apply to a wide range of shape and imaging datasets. In this paper, we use a tool from integral geometry and differential topology called the Euler characteristic (EC) transform [1–4], which maps shapes into vectors without requiring pre-specified landmark points or pairwise correspondences. This property is central to our innovations.

After finding a vector representation of the shape, our second challenge is quantifying which topological features are most relevant in explaining variation in a continuous outcome or binary class label. We address this classic take on variable selection by using a Bayesian regression model and an information theoretic metric to measure the relevance of each topological feature. Our Bayesian method allows us to perform variable selection for nonlinear functions — we discuss the importance of this requirement in Method Overview and Results.

The last challenge deals with how to interpret the most informative topological features obtained by our variable selection methodology. The EC transform is invertible; thus, we can take the most informative topological features and naturally recover the most imformative physical regions on the shape. In this paper, we introduce SINATRA: a unified statistical pipeline for sub-image analysis that addresses each of these challenges and is the first sub-image analysis method that does not require landmarks or correspondences.

Classically there have been three approaches to modeling random 3D images and shapes: *(i)* landmark-based representations [5,6], *(ii)* diffeomorphism-based representations [7], and *(iii)* representations that use integral geometry and excursions of random fields [8]. Landmark-based analysis uses points on shapes that are known to correspond with each other. As a result, any shape can be represented as a collection of 3D coordinates. Landmark-based approaches have two major shortcomings. First, many modern datasets are not defined by landmarks; instead, they consist of 3D CT scans [9,10]. Second, reducing these detailed mesh data to simple landmarks often results in a great deal of information loss.

Diffeomorphism-based approaches have bypassed the need for landmarks. Many tools have been developed that efficiently compare the similarity between shapes in large databases via algorithms that continuously deform one shape into another [11–16]. Unfortunately, these methods require diffeomorphisms between shapes: the map from shape *A* to shape *B* must be differentiable, as must the inverse of the map. Such functions are often called “correspondence maps” since they take two shapes and place them in correspondence. There are many applications with no such transformations because of qualitative differences. For example, in a dataset of fruit fly wings, some mutants may have extra lobes of veins [17]; or, in a dataset of brain arteries, many of the arteries cannot be continuously mapped to each other [18]. Indeed, in large databases such as the MorphoSource [10], the CT scans of skulls across many clades are not diffeomorphic. Recent algorithms have attempted to overcome this issue by constructing more general “functional” correspondences [19,20], which can be established even across shapes having different topology. However, there is a real need for 3D image analysis methods that do not require correspondences.

Previous work [2] introduced two topological transformations for shapes: the persistent homology (PH) transform and the EC transform. These tools from integral geometry first allowed for pairwise comparisons between shapes or images without requiring correspondence or landmarks. Since then, mathematical foundations of the two transforms and their relationship to the theory of sheaves and fiber bundles have been established [3, 4]. Detailed mathematical analyses have also been provided [3]. A nonlinear regression framework, which uses the EC transform to predict outcomes of disease free survival in glioblastoma [1], is most relevant to this paper. This works shows that the EC transform reduces the problem of regression with shape covariates into a problem in functional data analysis (FDA), and that nonlinear regression models are more accurate than linear models when predicting complex phenotypes and traits. The SINATRA pipeline further enhances the relation between FDA and topological transforms by enabling variable selection with shapes as covariates.

Beyond the pipeline, this paper includes software packaging to implement our approach and a detailed design of rigorous simulation studies which can assess the accuracy of sub-image selection methods. The freely available software comes with several built-in capabilities that are integral to sub-image analyses in both biomedical studies and geometric morphometric applications. First, SINATRA does not require landmarks or correspondences in the data. This means that the algorithm can be implemented on both datasets with raw unaligned shapes, as well as those that have been axis-aligned during preprocessing (see Supplementary Material). Second, given any dataset of 3D images, SINATRA outputs evidence measures that highlight the physical regions on shapes that explain the greatest variation between two classes. In many applications, users may suspect *a priori* that certain landmarks have greater variation across groups of shapes (e.g., via the literature). To this end, SINATRA also provides *P*-values and Bayes factors that detail how likely any region is identified by chance [21].

In this paper, we describe each mathematical step of the SINATRA pipeline and demonstrate its power and utility via simulations. We also use a dataset of mandibular molars from four different genera of primates to show that our method has the ability to *(i)* further understanding of how landmarks vary across evolutionary scales in morphology and *(ii)* visually detail how known anatomical aberrations are associated to specific disease classes and/or case-control studies.

## Method Overview

The SINATRA pipeline implements four key steps (Fig. 1). First, SINATRA summarizes the geometry of 3D shapes (represented as triangular meshes) by a collection of vectors (or curves) that encode changes in their topology. Second, a nonlinear Gaussian process model, with the topological summaries as input, classifies the shapes. Third, an effect size analog and corresponding association metric is computed for each topological feature used in the classification model. These quantities provide evidence that a given topological feature is associated with a particular class. Fourth, the pipeline iteratively maps the topological features back onto the original shapes (in rank order according to their association measures) via a reconstruction algorithm. This highlights the physical (spatial) locations that best explain the variation between the two groups. Details of our implementation choices are detailed below, with theoretical support given in the Supplementary Material.

**Figure 1.**
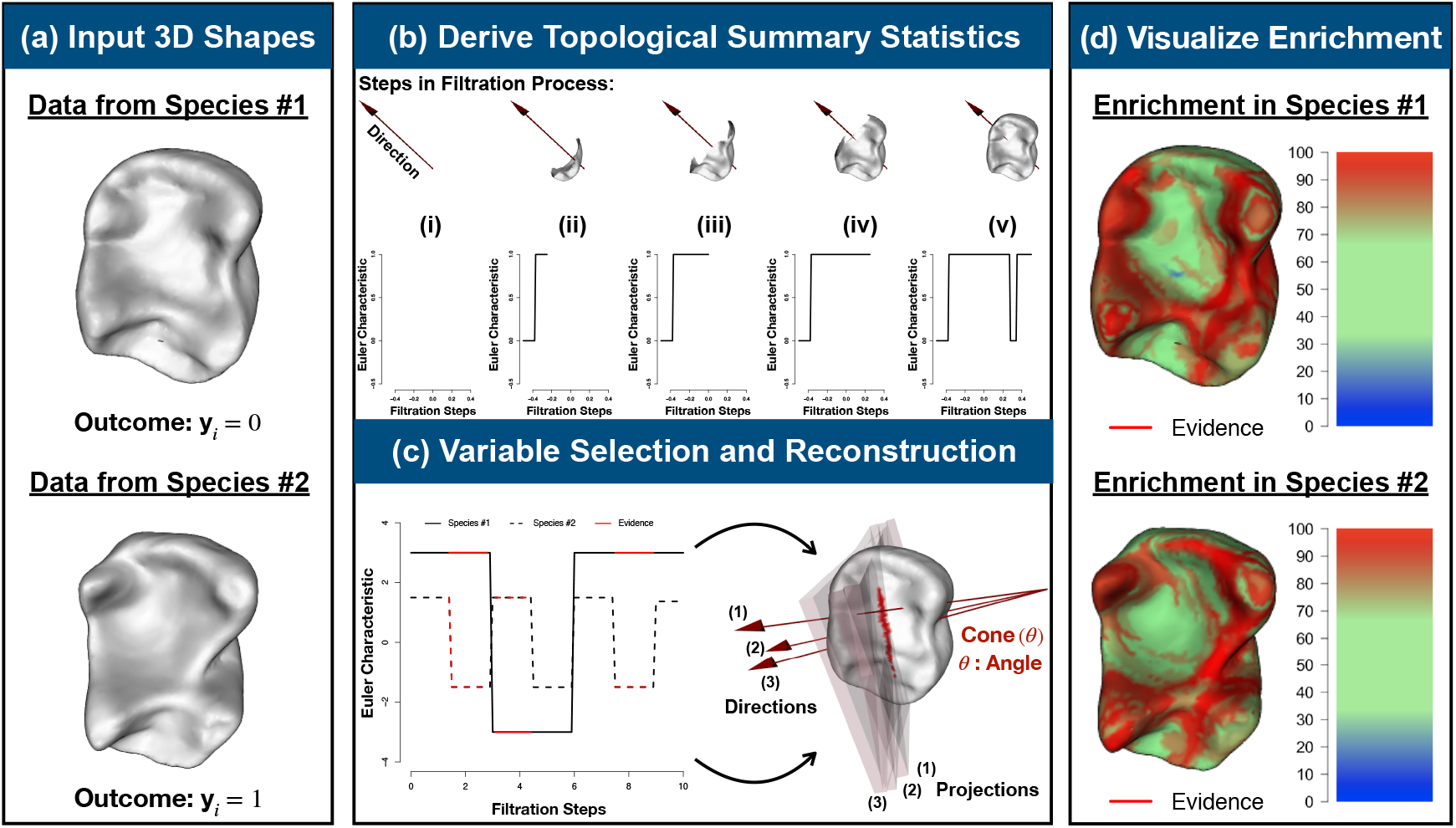
Schematic overview of SINATRA: a novel statistical framework for feature selection and association mapping with 3D shapes. **(a)** The SINATRA algorithm requires the following inputs: *(i)* aligned shapes represented as meshes; *(ii)* **y**, a binary vector denoting shape classes; *(iii) r*, the radius of the bounding sphere for the shapes; *(iv) c*, the number of cones of directions; *(v)* d, the number of directions within each cone; *(vi) θ*, the cap radius used to generate directions in a cone; and *(vii) l*, the number of sublevel sets (i.e., filtration steps) to compute the Euler characteristic (EC) along a given direction. Guidelines for how to choose the free parameters are given in Supplementary Table 1. **(b)** We select initial positions uniformly on a unit sphere. Then for each position, we generate a cone of *d* directions within angle *θ* using Rodrigues’ rotation formula [70], resulting in a total of m = *c* × *d* directions. For each direction, we compute EC curves with *l* sublevel sets. We concatenate the EC curves along all the directions for each shape to form vectors of topological features of length *p* = *l* × *m*. Thus, for a study with *n*-shapes, an *n* × *p* design matrix is statistically analyzed using a Gaussian process classification model. **(c)** Evidence of association for each topological feature vector are determined using relative centrality measures. We reconstruct corresponding shape regions by identifying the vertices (or locations) on the shape that correspond to “statistically associated” topological features. **(d)** This enables us to visualize the enrichment of physical features that best explain the variance between the two classes. The heatmaps display vertex evidence potential on a scale from [0 – 100]. A maximum of 100 represents the threshold at which the first shape vertex is reconstructed, while 0 denotes the threshold when the last vertex is reconstructed.

### Topological Summary Statistics for 3D Shapes

In the first step of the SINATRA pipeline, we use a tool from integral geometry and differential topology called the Euler characteristic (EC) transform [1–4]. For a mesh 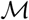, the Euler characteristic is an accessible topological invariants derived from:

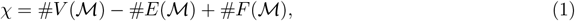

where 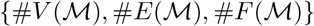 denote the number of vertices (corners), edges, and faces of the mesh, respectively. An EC curve 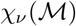 tracks the change in the Euler characteristic with respect to a given filtration of length *l* in direction *ν* (Figs. 1(a) and (b)). Theoretically, we first specify a height function 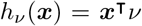 for vertex 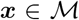 in direction *v*. We then use this height function to define sublevel sets (or subparts) of the mesh 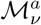 in direction *v*, where *h_ν_*(*x*) ≤ *a*. In practice, the EC curve is 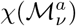 computed over a range of *l* filtration steps in direction *v* (Fig. 1(b)).

The EC transform is the collection of EC curves across a set of directions *v* = 1,…, *m*, and maps a 3D shape into a concatenated *p* = (*l × m*)-dimensional feature vector. For a study with *n*-shapes, an *n × p* design matrix **X** is statistically analyzed, where the columns denote the Euler characteristic computed at a given filtration step and direction. Each sublevel set value, direction, and set of shape vertices used to compute an EC curve are stored for the association mapping and projection phases of the pipeline. Previously, Curry et al. proved sufficiency, stating that theoretical upper bound on the minimum number of directions m required for the EC transform to preserve all information for a family of 3D shapes (see Theorem 7.14 in [3]) is estimated by

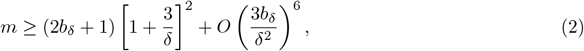

where *δ* is a lower bound on the “curvature” at every vertex on the mesh and *b_δ_* is a uniform upper bound on the number of critical values for the Euler characteristic curves when viewed in any given δ-ball of directions. While this upper bound may not yield optimal results in practice, we do use this theory to guide the collection of topological statistics with the general notion that considering larger values of *m* will lead to more robust summarization of shape variation. In this paper, we use a series of simulations and sensitivity analyses to outline empirical trends and develop intuition behind practically choosing the number of directions m and setting the granularity of sublevel filtrations *l* for real data.

### Statistical Model for Shape Classification

In the second step of the SINATRA pipeline, we use (weight-space) Gaussian process probit regression to classify shapes based on their topological summaries generated by the EC transformation. Namely, we specify the following (Bayesian) hierarchical model [22–26]

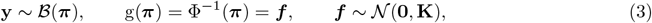

where **y** is an *n*-dimensional vector of Bernoulli distributed class labels, **π** is an *n*-dimensional vector representing the underlying probability that a shape is classified as a “case” (i.e., *y* = 1), *g*(·) is a probit link function with Φ(·) the cumulative distribution function (CDF) of the standard normal distribution, and ***f*** is an *n*-dimensional vector estimated from the data.

The key objective of SINATRA is to use the topological features in **X** to find the physical 3D properties that best explain the variation across shape classes. To do so, we use kernel regression, where the utility of generalized nonparametric statistical models is well-established due their ability to account for various complex data structures [27–32]. Generally, kernel methods posit that ***f*** lives within a reproducing kernel Hilbert space (RKHS) defined by some (nonlinear) covariance function, which implicitly account for higher-order interactions between features, leading to more complete classifications of data [33–35]. To this end, we assume ***f*** is normally distributed with mean vector **0**, and covariance matrix **K** defined by the radial basis function **K**_*ij*_ = exp{−*θ*||**x**_*i*_ − **x**_*j*_||^2^} with bandwidth *θ* set using the median heuristic to maintain numerical stability and avoid additional computational costs [36]. The full model specified in Equation (3) is commonly referred to as “Gaussian process classification” or GPC.

### Interpretable Feature (Variable) Selection

To estimate the model in Equation (3), we use an elliptical slice sampling Markov chain Monte Carlo (MCMC) algorithm (Supplementary Material, Section 1.1). Since we take a “weight-space” view on Gaussian processes, the model fitting procedure scales with the number of 3D meshes in the data, rather than with the number of topological features. The MCMC algorithm allows samples from the approximate posterior distribution of ***f*** (given the data), and also allows for the computation of an effect size analog for each topological summary statistic [37–39]

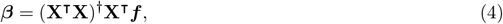

where 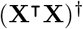 is the generalized inverse of 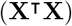.

These effect sizes represent the nonparametric equivalent to coefficients in linear regression using generalized least squares. SINATRA uses these weights and assigns a measure of relative centrality to each summary statistic (first panel Fig. 1(c)) [39]. This criterion evaluates how much information in classifying each shape is lost when a particular topological feature is removed from the model. This loss is determined by computing the Kullback-Leibler divergence (KLD) between *(i)* the conditional posterior distribution *p*(*β_−j_* | *β_j_* = 0) with the effect of the *j*-th topological feature set to zero, and *(ii)* the marginal posterior distribution *p*(*β_−j_*) with the effects of the *j*-th feature integrated out:

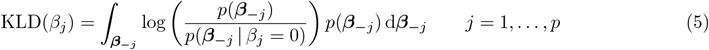

which has a closed form solution when the posterior distribution of the effect sizes is assumed to be (approximately) Gaussian (Supplementary Material, Section 1.2). Finally, we normalize to obtain an association metric for each topological feature, γ_*j*_ = KLD(*β_j_*)/∑ KLD(*β_j_*).

There are two main takeaways from this formulation. First, the KLD is non-negative, and equals zero if and only if the posterior distribution of *β_−j_* is independent of the effect *β_j_*. Intuitively, this says that removing an unimportant shape feature has no impact on explaining the variance between shape classes. Second, γ is bounded on the unit interval [0,1] with the natural interpretation of providing relative evidence of association for shape features; higher values suggest greater importance. For this metric, the null hypothesis assumes that every feature equally contributes to the total variance between shape classes, while the alternative proposes that some features are more central than others [39]. As we show in the Results and Supplementary Material, when the null assumption is met, SINATRA displays association results that appear uniformly distributed and effectively indistinguishable.

### Shape Reconstruction

After obtaining association measures for each topological feature, we map this information back onto the physical shape (second panel Fig. 1(c) and 1(d)). We refer to this process as *reconstruction*, as this procedure recovers regions that explain the most variation between shape classes (Supplementary Material, Section 1.3). Intuitively, we want to identify vertices on the shape that correspond to the topological features with the greatest association measures.

Begin by considering *d* directions within a cone of cap radius or angle *θ*, which we denote as 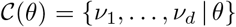. Next, let 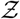 be the set of vertices whose projections onto the directions in 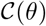 are contained within the collection of “significant” topological features — for every 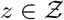, the product *z · v* is contained within a sublevel set (taken in the direction 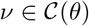) that shows high evidence of association in the feature selection step.

A reconstructed region is then defined as the union of all mapped vertices from each cone, or 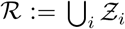. We use cones because vectors of Euler characteristics taken along directions close together express comparable information. That similiarity lets us leverage findings between them to increase our power of detecting truly associated shape vertices and regions — as opposed to antipodal directions where the lack of shared information may do harm when determining reconstructed manifolds (Supplementary Material, Section 1.4) [3,40,41].

### Visualization of Enriched Shape Regions

Once shapes have been reconstructed, we can visualize the relative importance or “evidence potential” for each vertex on the mesh with a simple procedure. First, we sort the topological features from largest to smallest according to their association measures γ_1_ ≥ γ_2_ ≥ ··· ≥ γ_*p*_. Next, we iterate through the sorted measures *T_k_* = (starting with *k* =1), and reconstruct the vertices corresponding to the topological features in the set {*j*: γ_*j*_ ≥ *T_k_*}.

The evidence potential for each vertex is defined as the largest threshold *T_k_* at which it is reconstructed for the first time, because vertices with earlier “birth times” in the reconstruction are more important relative to vertices that appear later. We illustrate these values via heatmaps over the reconstructed meshes (Fig. 1(d)). For consistency across different applications and case studies, we set the coloring of these heatmaps on a scale from [0 – 100]. A maximum value of 100 represents the threshold value at which the first vertex is born, while 0 denotes the threshold when the last vertex on the shape is reconstructed. Under the null hypothesis, where there are no meaningful regions differentiating between two classes of shapes, (mostly) all vertices appear to be born relatively early and at the same time (Supplementary Fig. 4). This is not the case under the alternative.

### Algorithm and Implementation

To facilitate analyses, software for implementing the SINATRA pipeline is carried out in `R` code and is freely available at https://github.com/lcrawlab/SINATRA. This algorithm requires these inputs:

- shapes represented as meshes (unaligned or preprocessed and axis-aligned);
- **y**, a binary vector denoting shape classes;
- *r*, the radius of the bounding sphere for the shapes (which we usually set to 1/2 since we work with meshes normalized to the unit ball);
- *c*, the number of cones of directions;
- *d*, the number of directions within each cone;
- *θ*, the cap radius used to generate directions in a cone;
- *l*, the number of sublevel sets (i.e., filtration steps) to compute the Euler characteristic (EC) along a given direction.

A table with general guidelines for how set these free parameters in practice is given in Supplementary Table 1. In the next section, we discuss strategies for choosing values for the free parameters through simulation studies. A table detailing the scalability for the current algorithmic implementation of SINATRA can also be found in the Supplementary Material (see Supplementary Table 2). For some context, runtime for SINATRA is dependent upon both the number of meshes in a given study (*n*), as well as the number of cones of directions (*c*), directions within each cone (*d*), and sublevel sets (*l*) used to compute the EC curves. Note that the latter three parameters all contribute to the total number of topological statistics used summarize each mesh (i.e., *p* = *l* × *m* EC features taken over *m* = *c* × *d* total directions). Overall, the SINATRA algorithm scales (approximately) linearly in the number of shapes. For example, while holding the values of {*c* = 25, *d* = 5, *l* = 25} constant, it takes ~60 and ~96 seconds to analyze datasets with *n* = 25 and *n* = 100 shapes, respectively. The rate limiting step in the SINATRA pipeline is the computation of the association metric for each topological summary statistic (see Equation (5)). To mitigate this burden, we use approximations such that the cost of calculating each *γ_j_* is made up of *p*-independent *O*(*p*^2^) operations, which can be parallelized (see derivations provided in Supplementary Material, Section 1.2).

## Results

### Simulation Study: Perturbed Spheres

We begin with a proof-of-concept simulation study to demonstrate both the power of our proposed pipeline and how different parameter value choices affect its ability to detect associated features on 3D shapes. Again, a table with general guidelines for how set the free parameters within the SINATRA pipeline can be found in Supplementary Table 1. Here, we take 100 spheres and perturb regions, or collections of vertices, on their surfaces to create two classes with a two-step procedure:

1. We generate a fixed number of (approximately) equidistributed regions on each sphere: some number *u* regions to be shared across classes, and the remaining v regions to be unique to class assignment.
2. To create each region, we perturb the *k* closest vertices {***x***_1_, ***x***_2_,…, ***x**_k_*} by a pre-specified scale factor *α* and add some random normally distributed noise 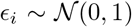 by setting 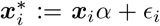 for *i* = 1, …, *k*.

We consider three scenarios based on the number of shared and unique regions between shape classes (Figs. 2(a)–2(c)). We choose (*u, v*) = (2,1) (scenario I), (6, 3) (scenario II), and (10, 5) (scenario III), and set all regions to be *k* = 10 vertices.

**Figure 2.**
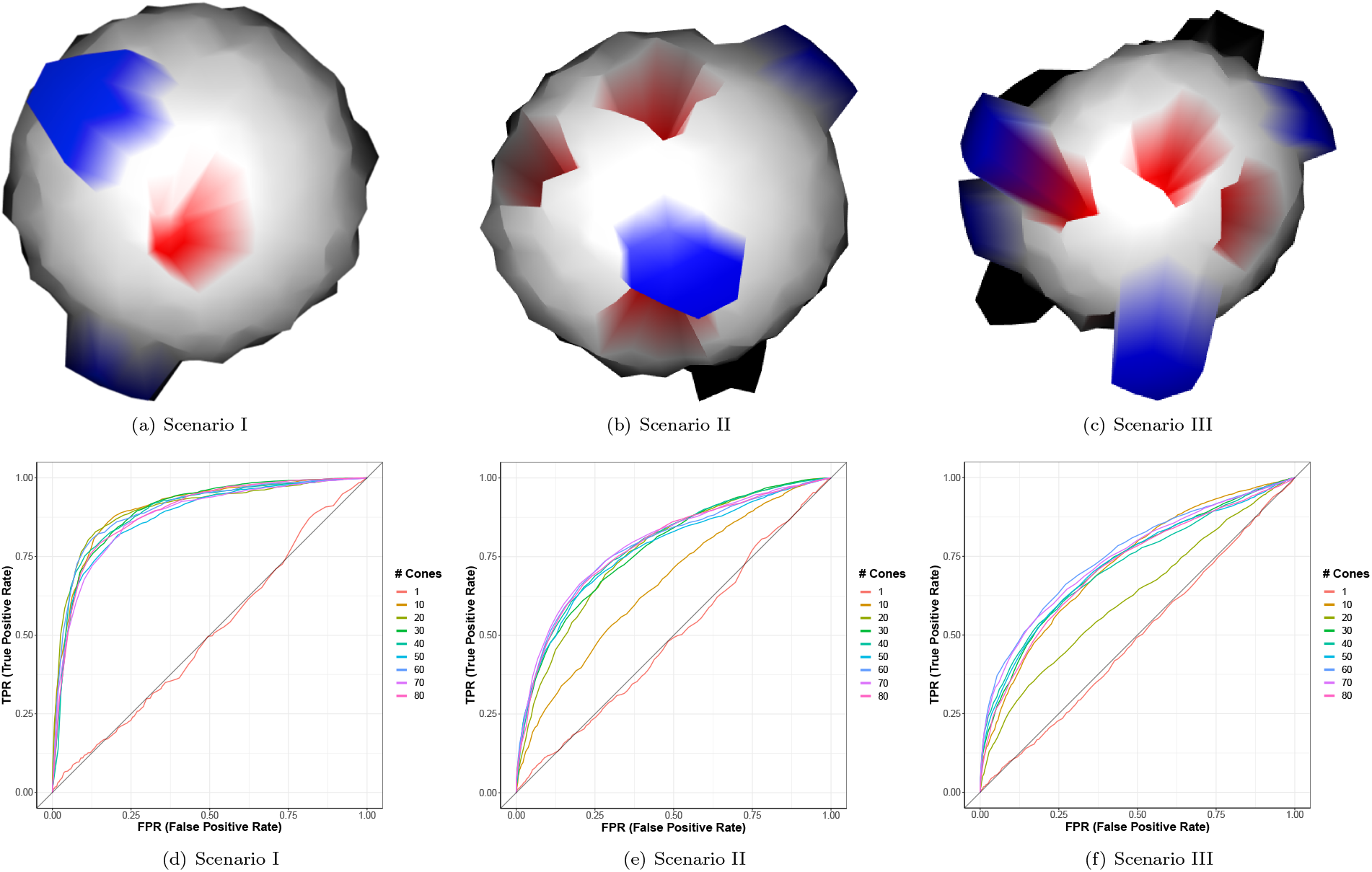
Power and sensitivity analysis for detecting associated vertices across different classes of perturbed spheres. We generate 100 shapes by partitioning unit spheres into regions 10 vertices-wide, centered at 50 equidistributed points. Two classes (50 shapes per class) are defined by shared (blue protrusions) and class-specific (red indentations) characteristics. The shared or “non-associated” features are chosen by randomly selecting *u* regions and pushing the sphere outward at each of these positions. To generate class-specific or “associated” features, *v* distinct regions are chosen for a given class and perturbed inward. We vary these parameters and analyze three increasingly more difficult simulation scenarios: **(a)** *u* = 2 shared and *v* =1 associated; **(b)** *u* = 6 shared and *v* = 3 associated; and **(c)** *u* = 10 shared and *v* = 5 associated. In panels **(d)**-**(f)**, ROC curves depict the ability of SINATRA to identify vertices located within associated regions, as a function of increasing the number of cones of directions used in the algorithm. These results give empirical evidence that seeing more of a shape (i.e., using more unique directions) generally leads to an improved ability to map back onto associated regions. Other SINATRA parameters were fixed: *d* = 5 directions per cone, *θ* = 0.15 cap radius used to generate directions in a cone, and *l* = 30 sublevel sets per filtration. Results are based on 50 replicates in each scenario.

Each sequential scenario represents an increase in degree of difficulty, because class-specific regions should be harder to identify in shapes with more complex structures. We analyze 50 unique simulated datasets for each scenario. In each dataset, only the *v*-region vertices used to create class-specific regions are defined as true positives, and we quantify SINATRA’s ability to prioritize these true vertices using receiver operating characteristic (ROC) curves plotting true positive rates (TPR) against false positive rates (FPR) (Supplementary Material, Section 2). We then evaluate SINATRA’s power as a function of its free parameter inputs: *c* number of cones, *d* number of directions per cone, direction generating cap radius *θ*, and *l* number of sublevel sets per filtration. We iteratively vary each parameter, while holding the others as constants {*c* = 25, *d* = 5, *θ* = 0.15, *l* = 30}. Figures displayed in the main text are based on varying the number of cones (Figs. 2(d)–2(f)), while results for the other sensitivity analyses can be found in the Supplementary Material (Supplementary Figs. 1-3).

As expected, SINATRA’s performance is consistently better when shapes are defined by a few prominent regions (e.g., scenario I) versus when shape definitions are more complex (e.g., scenarios II and III), because each associated vertex makes a greater individual contribution to the overall variance between classes (i.e., 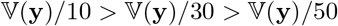). Similar trends in performance have been shown during the assessment of high-dimensional variable selection methods in other application areas [42–44].

This simulation study also demonstrates the general behavior and effectiveness of the SINATRA algorithm as a function of different choices for its free input parameters. First, we assess how adjusting the number of cones of directions used to compute Euler characteristic curves changes power. Computing topological summary statistics over just a single cone of directions (i.e., *c* = 1) is ineffective at capturing enough variation to identify class-specific regions (Figs. 2(d)–2(f)), which supports the intuition that seeing more of a shape leads to an improved ability to understand its complete structure [1–3]. Our empirical results show that more power can be achieved by summarizing the shapes with filtrations taken over multiple directions. In practice, we suggest specifying multiple cones *c* > 1 and utilizing multiple directions *d* per cone (see monotonically increasing power in Supplementary Fig. 1).

While the other two parameters (*θ* and l) do not have monotonic properties, their effects on SINATRA’s performance still have natural interpretations. For example, when changing the angle between directions within cones from *θ* ∈ [0.05, 0.5] radians, we observe that power steadily increases until *θ* = 0.25 radians and then slowly decreases afterwards (Supplementary Fig. 2). This supports previous theoretical results that cones should be defined by directions in close proximity to each other [3]; but not so close that they explain the same local information with little variation.

Perhaps most importantly, we must understand how the number of sublevel sets *l* (i.e., the number of steps in the filtration) used to compute Euler characteristic curves affects the performance of the algorithm. As we show in the next section, this function depends on the types of shapes being analyzed. Intuitively, for very intricate shapes, coarse filtrations with too few sublevel sets cause the algorithm to miss or “step over” very local undulations in a shape. For the spheres simulated in this section, class-defining regions are global-like features, and so finer filtration steps fail to capture broader differences between shapes (Supplementary Fig. 3); however, this failure is less important when only a few features decide how shapes are defined (e.g., scenario I). In practice, we recommend choosing the angle between directions within cones *θ* and the number of sublevel sets *l* via cross validation or some grid-based search.

As a final demonstration, we show what happens when we meet the null assumptions of the SINATRA pipeline (Supplementary Fig. 4). Under the null hypothesis, our feature selection measure assumes that all 3D regions of a shape equally contribute to explaining the variance between classes — that is, no one vertex (or corresponding topological characteristics) is more important than the others. We generate synthetic shapes under the two cases when SINATRA fails to produce significant results: (a) two classes of shapes that are effectively the same (up to some small Gaussian noise), and (b) two classes of shapes that are completely dissimilar. In the first simulation case, there are no “significantly associated” regions and thus no group of vertices stand out as important (Supplementary Fig. 4(a)). In the latter simulation case, shapes between the two classes look nothing alike; therefore, all vertices contribute to class definition, but no one feature is key to explaining the observed variation (Supplementary Fig. 4(b)).

### Simulation Study: Caricatured Shapes

Our second simulation study modifies computed tomography (CT) scans of real Lemuridae teeth (one of the five families of Strepsirrhini primates commonly known as lemurs) [11] using a well-known caricaturization procedure [45]. We fix the triangular mesh of an individual tooth and specify class-specific regions centered around known biological landmarks (Fig. 3) [11]. For each triangular face contained within a class-specific region, we apply a corresponding affine transformation, positively scaled, that smoothly varies on the triangular mesh and attains its maximum at the biological landmark used to define the region (Supplementary Material, Section 3). We caricature 50 different teeth with two steps (Fig. 3(a)):

1. Assign *v* of a given tooth’s landmarks to be specific to one class and *v′* to be specific to the other class.
2. Perform the caricaturization: multiply each face in the *v* and *v′* class-specific regions by a positive scalar (i.e., exaggerated or enhanced). Repeat twenty-five times (with some small noise per replicate) to create two equally-sized classes of 25 shapes.

**Figure 3.**
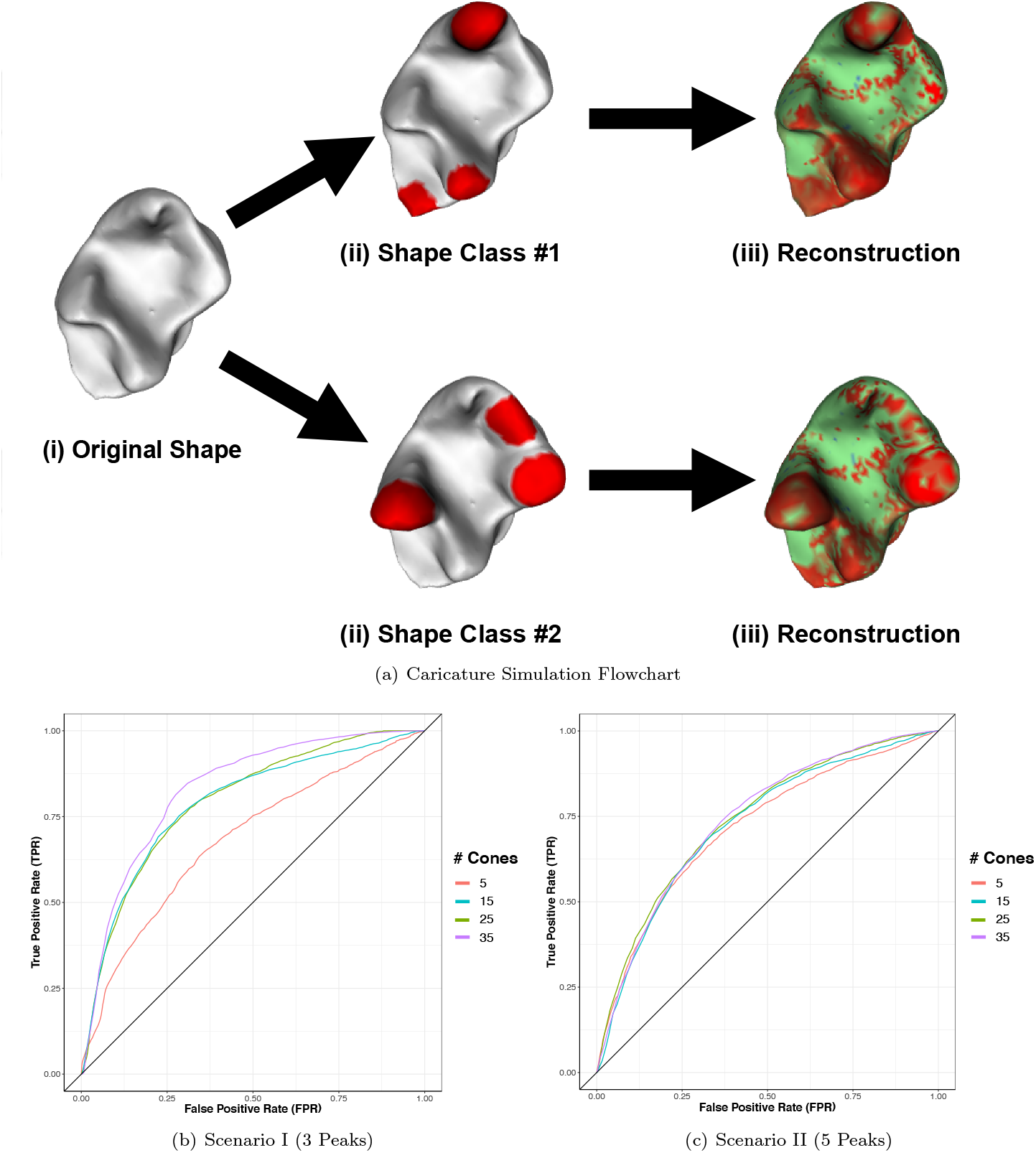
Power and sensitivity analysis for detecting associated vertices across different classes of caricatured shapes. **(a)** We modify real Lemuridae molars using the following caricaturization procedure: *(i)* fix the triangular mesh of an individual tooth; *(ii)* assign *v* of the known landmarks for the tooth [11] to be specific to one class and *v′* to be specific to the other. The caricaturization is performed by positively scaling each face within these regions so that class-specific features are exaggerated. We repeat twenty-five times (with some small added noise) to create two classes of 25 shapes. *(iii)* The synthetic shapes are analyzed by SINATRA to identify the associated regions. We consider two scenarios by varying the number of class-specific landmarks that determine the caricaturization in each class. In scenario I, we set *v,v′* = 3; and in scenario II, *v,v′* = 5. In panels **(b)** and **(c)**, ROC curves depict the ability of SINATRA to identify vertices located within associated regions, as a function of the number of cones of directions used in the algorithm. Other SINATRA parameters were fixed: *d* = 5 directions per cone, *θ* = 0.15 cap radius used to generate directions in a cone, and *l* = 50 sublevel sets per filtration. Results are based on 50 replicates in each scenario.

We explore two scenarios by varying the number of class-specific landmarks *v* and *v′* that determine the caricaturization in each class. First we set both *v*, *v′* = 3; next, we fix *v*, *v′* = 5. Like the simulations with perturbed spheres, the difficulty of the scenarios increases with the number of caricatured regions. We evaluate SINATRA’s ability to identify the vertices involved in the caricaturization using ROC curves (Supplementary Material, Section 2), and we assess this estimate of power as a function of the algorithm’s free parameter inputs. While varying each parameter, we hold the others as constants {*c* = 15, *d* = 5, *θ* = 0.15, *l* = 50}. Figures in the main text are based on varying the number of cones *c* (Figs. 3(b) and 3(c)); results for the other sensitivity analyses can be found in the Supplementary Material (Supplementary Figs. 5-7).

Overall, using fewer caricatured regions results in better (or at least comparable) performance. Like the simulations with perturbed spheres, SINATRA’s power increases monotonically with an increasing number of cones and directions used to compute the topological summary statistics (Figs. 3(b), 3(c), and Supplementary 5). For example, at a 10% FPR with *c* = 5 cones, we achieve 30% TPR in scenario I experiments and 35% TPR in scenario II. Increasing the number of cones to *c* = 35 improves power to 52% and 40% TPR for scenarios I and II, respectively. Trends from the previous section continue when choosing the angle between directions within cones (Supplementary Fig. 6) and the number of sublevel sets (Supplementary Fig. 7). Results for the perturbed spheres suggest that there is an optimal cap radius for generating directions in a cone. Since we are analyzing shapes with more intricate features, finer filtrations lead to more power.

### Simulation Study: Method Comparisons

In this subsection, we compare the power of SINATRA to other state-of-the-art sub-image selection methods. Here, we revisit the same simulation scenarios using the perturbed spheres and caricatured teeth simulation schemes, respectively. Once again, we use ROC curves plotting TPR versus FPR to assess the ability of each method to identify vertices in class-specific associated regions. For SINATRA, we use a grid search to choose the algorithm’s free parameters. This led to us setting {*c* = 60, *d* = 5, *θ* = 0.15, *l* = 30} for the perturbed spheres and {*c* = 35, *d* = 5, *θ* = 0.15, *l* = 50} for the caricatured teeth. Note that these final values follow the trends we observed from the sensitivity analyses presented in the previous two subsections. We then consider the following three types of competing approaches:

- **Vertex-Level Regularization (Baseline):** Using the fact that all the vertices within a set of simulated meshes are in one-to-one correspondence, we vectorize the shapes by concatenating the (*x, y, z*)-coordinates of each vertex into a single vector. In this way, we have a (3 × *q*)-dimensional vector representation of each shape in our dataset, where *q* is the number of vertices that each mesh contains. In other words, each vertex is directly represented by the three entries of the vector (once for each dimension of the shape). This results in a final *n* × (3 × *q*) design matrix, where *n* is the total number of shapes in the data. To perform sub-image selection, we implement Elastic Net regularization [46] using the `glmnet` package [47] in `R` (with the penalization term chosen via cross-validation) to assign sparse individual coefficients to each column of the design matrix. Power is then determined by either ranking the mean or maximum of the three coefficients corresponding to the (*x, y, z*)-coordinates for each vertex.
- **Landmark-Level Regularization:** We implement this method by taking advantage of the fact that the simulated objects in our study are generated from perturbations of a base shape. Here, we generate landmarks by selecting *v* = {500, 2000} equidistributed points on a base shape (in the sense of Euclidean distance). Next, we concatenate the (*x, y, z*)-coordinates of the *k* closest vertices to each point and obtain landmark-specific vectors in 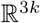. This results in a final *n* × (3 × *v* × *k*) design matrix, where again *n* is the total number of shapes in the data. To perform sub-image selection with this method, we use a logistic regression model with Group Lasso regularization [48], where the groups are predefined as the coordinates belonging to a given landmark and the group-based penalization term is chosen via cross-validation with the `gglasso` [49] in `R`. Note that we consider a Group Lasso penalty in order to encourage selection of important landmarks as a whole, rather than individual vertices. True and false positives are then determined by ranking each landmarks based on the magnitude of their coefficients.
- **Limit Shapes Algorithm [20]:** This algorithm builds upon a “functional map network” consisting of (dual representations of) point-by-point pairwise correspondences between all pairs of shapes in a dataset. From here, a consistent latent basis can be extracted and used for constructing a “limit shape” underlying the given collection of 3D objects but represented in a latent space. Deviations of individual shapes from this “limit shape,” represented as functions defined on these shapes, provide a visual guide highlighting sources of variability that often carry rich semantic meanings. In our simulations, we generate maps between all mesh pairs using a state-of-the-art pipeline that searches for the optimal “bounded-distortion map” and interpolates as many “Gaussian process landmarks” as possible [16]. These point-to-point maps of bounded conformal distortion are then converted into functional maps which are essentially the same maps but represented under a different system of bases of eigenfunctions of the discrete Laplace-Beltrami operators. The resulting functional maps are directly fed into the Limit Shapes workflow [20]. To perform sub-image selection, we compute a vertex importance vector using the first 10 eigenfunctions of the discrete Laplace-Beltrami operator corresponding to the 10 smallest eigenvalues. Vertex weights were summarized within the eigenfunctions by scaling the absolute value of each entry *ψ_i_* by the maximum vertex weight within that eigenfunction such that *W_j_* = |*ψ_j_*|/max{|*ψ*_1_|,…, |*ψ_n_*|}. After norming weights within the eigenfunctions, we obtain a final vector of vertex weights by taking the entry-wise maximum of each vertex across the 10 eigenfunctions. Using this procedure enables us to rank features from Limit Shapes and to generate ROC curves. To imitate realistic conditions where point-by-point maps are not known *a priori*, we also introduce a (misspecified) variation of Limit Shapes in which the functional map network is partially scrambled.

Figures 4 and 5 display the performance of SINATRA and each of the competing methods on the perturbed spheres and caricatured teeth, respectively, across 50 replicates in each simulation scenario. There are a few important takeaways from these comparisons. First, the Group Lasso landmark-based method consistently performs the worst among all approaches that we consider, regardless of the number of landmarks used. This is likely due to over-regularization where landmarks containing only a few vertices with nonzero effects are still be treated as non-associated with class variation. The vertex-level Elastic Net baselines perform the best under the perturbed sphere simulations, particularly when the variation between shape classes is sparsely driven by just a few associated regions (see Fig. 4). Furthermore, since this approach uses a simple regression, it scales linearly with the number of vertices on each shape. The Limit Shapes algorithm also had a quicker runtime than SINATRA, which took approximately 30 minutes to run per analysis (Supplementary Table 2). While the Limit Shapes approach performs generally well in each simulation scenario, its performance drops significantly when the functional mapping input into the algorithm is misspecified. This highlights an important and practical advantage of SINATRA which maintains its utility even when such user-specified point-by-point correspondences are unknown. Overall, the performance of SINATRA remains competitive in all settings, but clearly becomes the best approach in the caricatured teeth simulations where there is an increase in the overall complexity of the meshes that are analyzed (see Fig. 5). We hypothesize that the simple vertex-level regularization approaches struggle on the caricatured teeth because the coordinate-based representation becomes less effective at capturing the varying topology and geometry between shapes.

**Figure 4.**
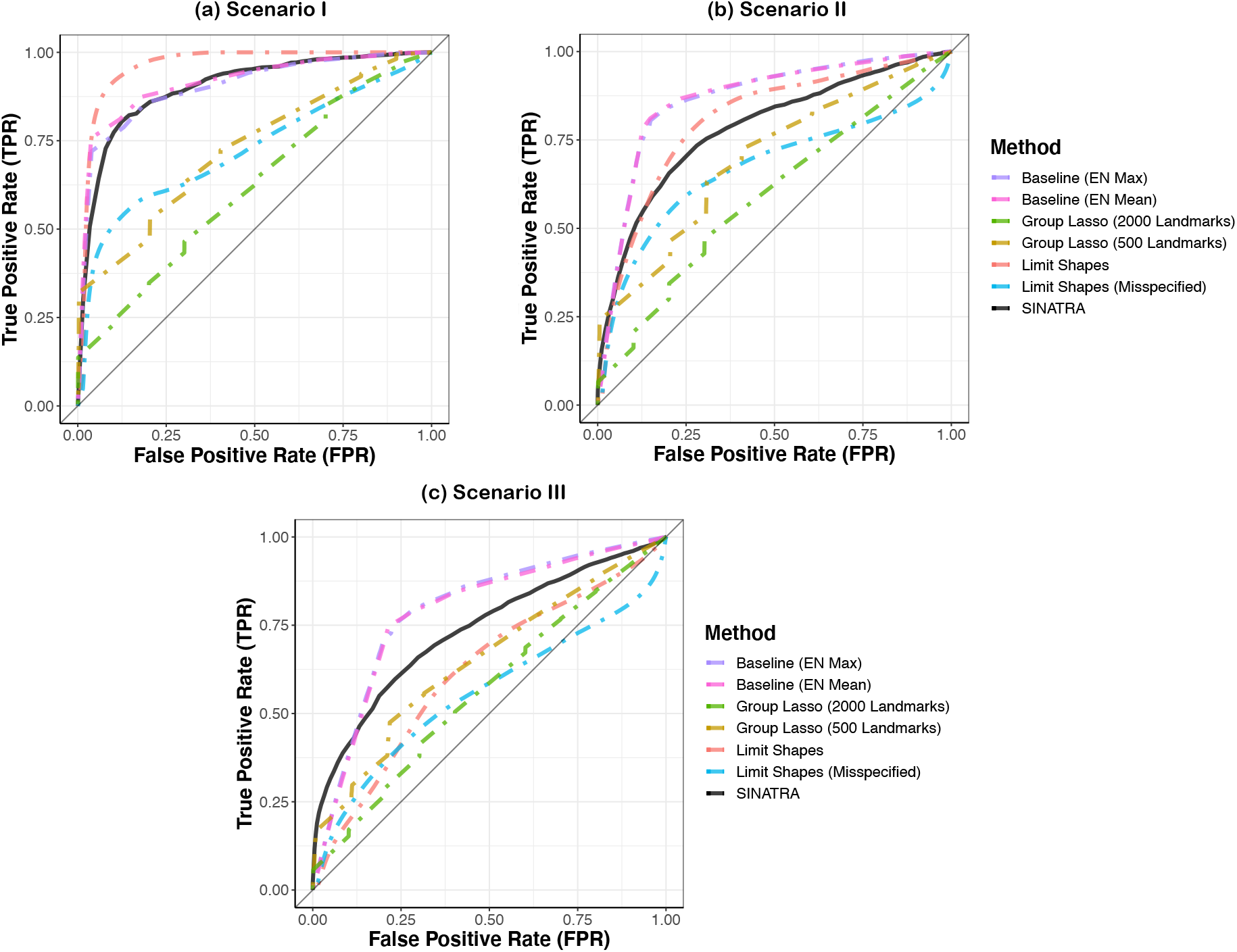
Receiver operating characteristic (ROC) curves comparing the performance of SINATRA with competing methods in the perturbed sphere simulations. Here, we generate 100 shapes by partitioning unit spheres into regions 10 vertex-wide, centered at 50 equidistributed points. Two classes (50 shapes per class) are defined by the number of shared *u* and class-specific *v* characteristics. We vary these parameters and analyze three increasingly more difficult simulation scenarios: **(a)** *u* = 2 shared and *v* = 1 associated; **(b)** *u* = 6 shared and *v* = 3 associated; and **(c)** *u* =10 shared and *v* = 5 associated. The ROC curves depict the ability of SINATRA to identify vertices located within associated regions using parameters {*c* = 60, *d* = 5, *θ* = 0.15, *l* = 30} chosen via a grid search. We compare SINATRA to three methods. The first baseline concatenates the (x, y, z)-coordinates of all vertices on the sphere and treats them as features in a data frame. It then uses Elastic Net regularization to assign sparse individual coefficients to each coordinate. For this method, we assess power by either taking the mean or maximum of the coefficient values corresponding to each vertex. The second method assumes 500 or 2000 equally spaced landmarks across each mesh and implements a Group Lasso penalty on the collection on vertices within these regions to rank associated features. Lastly, we compare the Limit Shapes algorithm [20] where normalized vertex weights are used to determine true and false positives. Note that the Limit Shapes algorithm requires a functional correspondence map between all pairs of meshes in the data; therefore, we display results for this algorithm both when a correspondence map is known and when the map has been misspecified. Results in each ROC curve are based on 50 replicates in each scenario.

**Figure 5.**
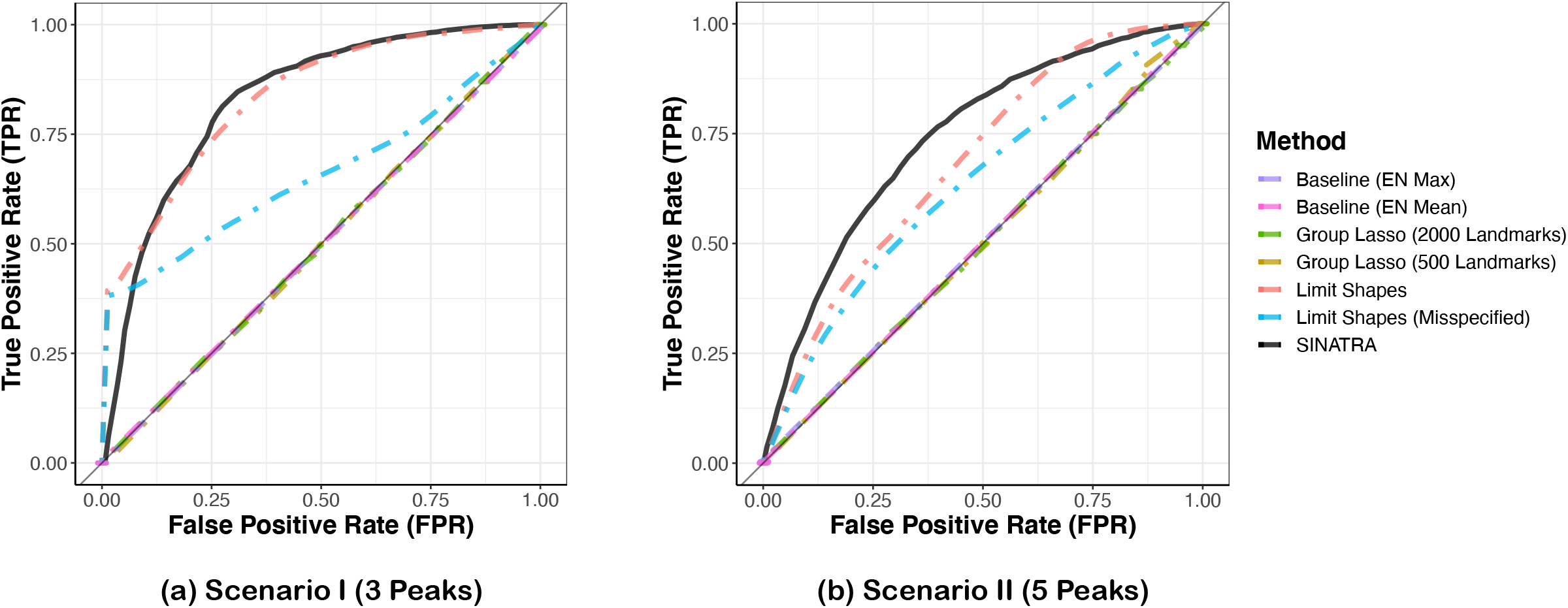
Receiver operating characteristic (ROC) curves comparing the performance of SINATRA with competing methods in the caricatured teeth simulations. Here, we modify real Lemuridae molars using a caricaturization procedure where we assign a *v* number of known landmarks for a tooth [11] to be specific to one class and *v′* to be specific to the other. We then consider two scenarios by varying the number of class-specific landmarks. In scenario I, we set *v*, *v′* = 3; and in scenario II, *v*, *v′* = 5. The ROC curves depict the ability of SINATRA to identify vertices located within associated regions using parameters {*c* = 35, *d* = 5, *θ* = 0.15, *l* = 50} chosen via a grid search. We compare SINATRA to three methods. The first baseline concatenates the (x, y, z)-coordinates of all vertices on each caricatured tooth and treats them as features in a data frame. It then uses Elastic Net regularization to assign sparse individual coefficients to each coordinate. For this method, we assess power by either taking the mean or maximum of the coefficient values corresponding to each vertex. The second method assumes 500 or 2000 equally spaced landmarks across each mesh and implements a Group Lasso penalty on the collection on vertices within these regions to rank associated features. Lastly, we compare the Limit Shapes algorithm [20] where normalized vertex weights are used to determine true and false positives. Note that the Limit Shapes algorithm requires a functional correspondence map between all pairs of meshes in the data; therefore, we display results for this algorithm both when a correspondence map is known and when the map has been misspecified. Results in each ROC curve are based on 50 replicates in each scenario.

### Recovering Known Morphological Variation Across Genera of Primates

As an application of our pipeline, with “ground truth” or known morphological variation, we consider a dataset of CT scans of *n* = 59 mandibular molars from two suborders of primates: Haplorhini (which include tarsiers and anthropoids) and Strepsirrhini (which include lemurs, galagos, and lorises). From the haplorhine suborder, 33 molars came from the genus *Tarsius* [11,50,51] and 9 molars from the genus *Saimiri* [52]. From the strepsirrhine suborder, 11 molars came from the genus *Microcebus* and 6 molars from the genus *Mirza* [11,50,51]; both are lemurs.

We chose this specific collection of molars because morphologists and evolutionary anthropologists understand variations of the paraconid, the cusp of a primitive lower molar. The paraconids are retained only by *Tarsius* and do not appear in the other genera (Fig. 6) [52, 53]. Using phylogenetic analyses of mitochondrial genomes across primates, Pozzie et al. estimate divergence dates of the subtree composed of *Microcebus* and *Mirza* from *Tarsius* at 5 million years before the branching of *Tarsius* from *Saimiri* [54]. We want to see if SINATRA recovers the information that the paraconids are specific to the *Tarsius* genus. We also investigate if variation across the molar is associated to the divergence time of the genera.

**Figure 6.**
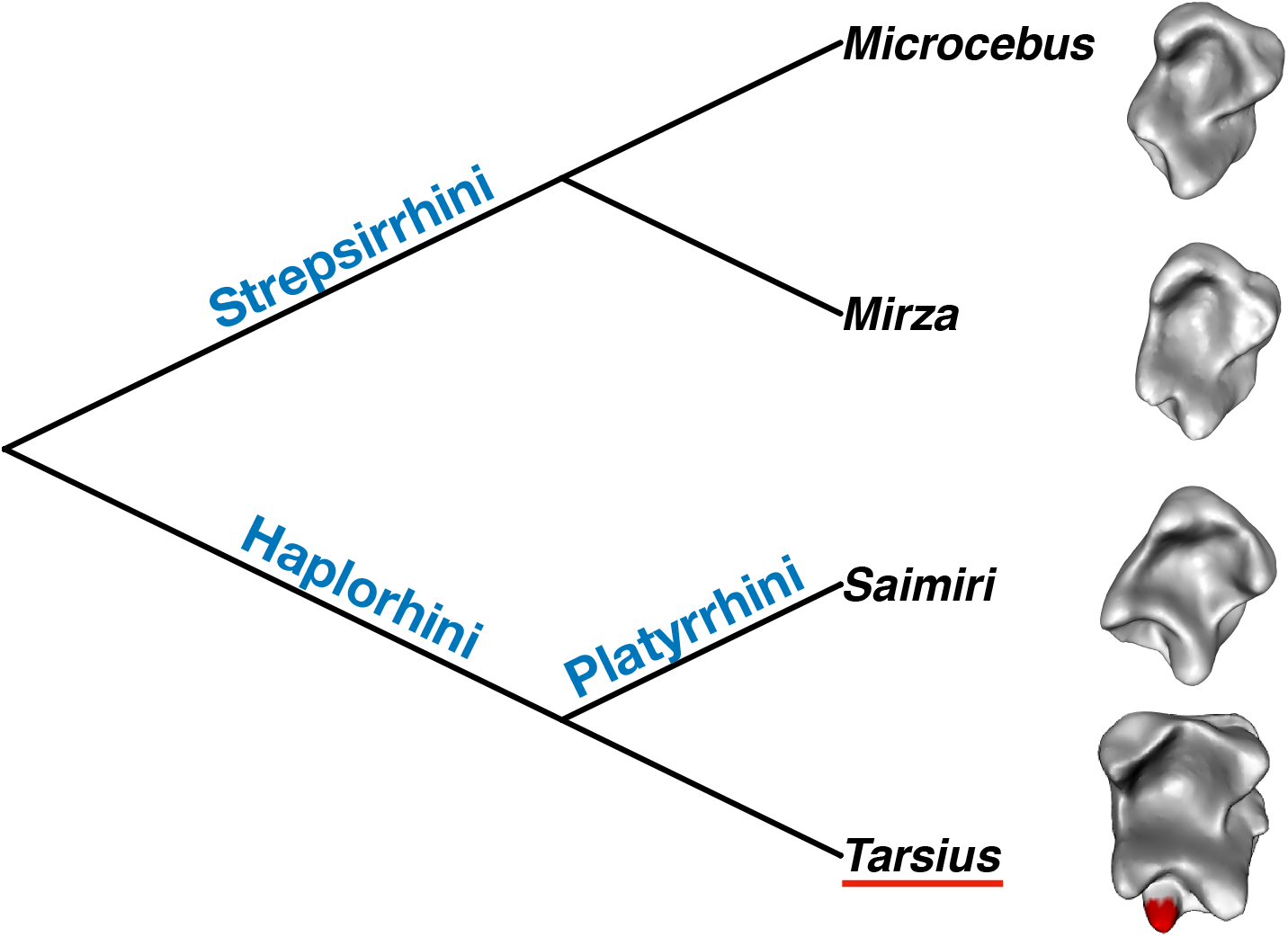
Illustration of the relationship between the phylogenetics and unique paraconids in molars belonging to primates in *Tarsius* genus. We carry out three pairwise comparisons to analyze the physical differences between *Tarsius* molars and teeth from *(i) Saimiri, (ii) Mirza*, and *(iii) Microcebus* genus. Here, we depict the phylogentic relationship between these groups. Morphologically, we know that tarsier teeth have an additional high-cusp (highlighted in red), which allows this genus of primate to eat a wider range of foods [71]. With these data, we want to assess SINATRA’s ability to find this region of interest (ROI).

For these analyses, we consider two types of data preprocessing procedures. In the first, the meshes of all teeth were pre-aligned, centered at the origin, and normalized within a unit sphere using the `auto3dgm` software [55] (Supplementary Fig. 8). In the second, we first conduct the Euler characteristic transformation on the raw data for each tooth and then we *implicitly* normalize the 3D meshes by aligning their EC curves (Supplementary Figs. 9 and 10). This latter analysis is used to demonstrate the effectiveness of SINATRA in settings where *a priori* correspondences between shapes are unavailable (details in Supplementary Material, Section 4). Since *Tarsius* is the only genus with the paraconid in this sample, we use SINATRA to perform three pairwise classification comparisons (*Tarsius* against *Saimiri*, *Mirza*, and *Microcebus*, respectively), and assess SINATRA’s ability to prioritize/detect the location of the paraconid as the region of interest (ROI). Based on our simulation studies, we run SINATRA with *c* = 35 cones, *d* = 5 directions per cone, a cap radius of *θ* = 0.25 to generate each direction, and *l* = 75 sublevel sets to compute topological summary statistics. In each comparison, we evaluate the evidence for each vertex based on the first time that it appears in the reconstruction: this is the evidence potential for a vertex. A heatmap for each tooth provides visualization of the physical regions that are most distinctive between the genera (Fig. 7). In this figure, we also display results from implementing the Limit Shapes algorithm [20] on the `auto3dgm` pre-aligned data as a baseline. Once again, for this method, we use the normalized vector weights to determine the importance of each vertex in the data.

**Figure 7.**
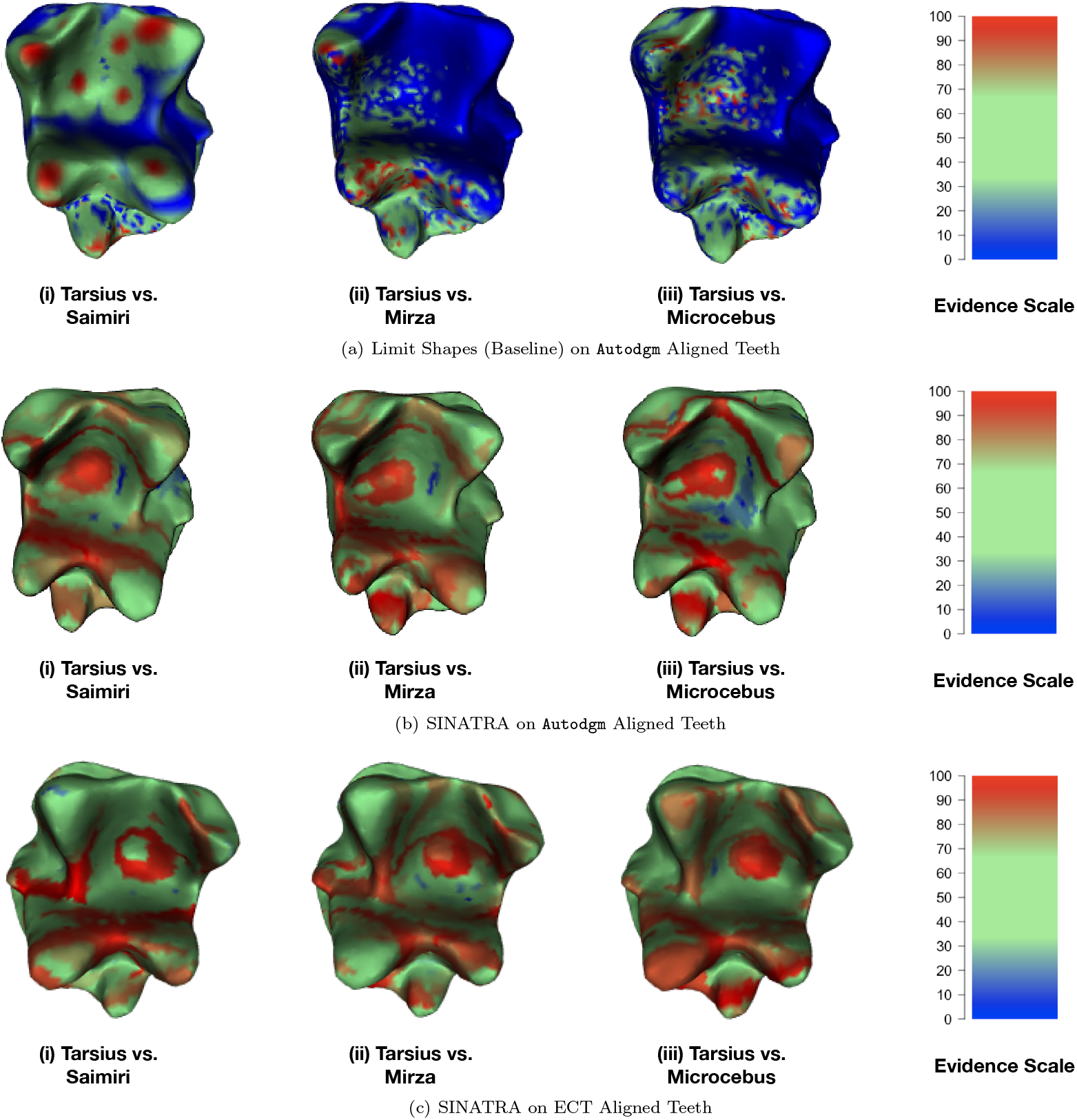
Real data analysis aimed at detecting unique paraconids in molars belonging to primates in the *Tarsius* genus. We carry out three pairwise comparisons to analyze the physical differences between *Tarsius* molars and teeth from *(i) Saimiri, (ii) Mirza*, and *(iii) Microcebus* genus. Here, we consider the region of interest (ROI) to be the additional high-cusp in the *Tarsius* molars (highlighted in red in Figure 6). In this figure, we give an example of the mesh reconstruction from each class comparison using the Limit Shapes algorithm [20] and SINATRA. Panels **(a)** and **(b)** show results on shapes that were pre-aligned with the `auto3dgm` software [55]. Panel **(c)** illustrates results using SINATRA without any prior knowledge of pairwise correspondence maps between shapes in the data. In this experiment, we first conduct the Euler characteristic transformation on the raw data for each tooth. Then we implicitly normalize the meshes by aligning the EC curves via the approximate grid search procedure detailed in Section 4 of the Supplementary Material. Overall, results are consistent with the phylogeny of the primates, as well as with our previous simulation studies. Genetically, *Tarsius* differ more from the *Mirza* and *Microcebus* genera, rather than from *Saimiri*. As a result, SINATRA finds the unique paraconid in the former two comparisons because of the appropriate genetic distance, rather than in the latter case where molar structures are more similar. This trend occurs whether or not we have pre-aligned data. The heatmaps in each panel display vertex evidence potential on a scale from [0 – 100]. A maximum of 100 represents the threshold at which the first shape vertex is reconstructed, while 0 denotes the threshold when the last vertex is reconstructed.

To assess the ability of the Limit Shapes and SINATRA to find *Tarsius*-specific paraconids, we use a null-based scoring method. We place a paraconid landmark on each *Tarsius* tooth, and consider the *K* = {10, 50, 100, 150, 200} nearest vertices surrounding the landmark’s centermost vertex. This collection of *K* +1 vertices defines our ROI. Within each ROI, we weight the Limit Shape or SINATRA-computed evidence potentials by the surface area (or area of the Voronoi cell) encompassed by their corresponding vertices, and then sum the scaled potentials together across the ROI vertices. This aggregated value, which we denote as *τ*\*, represents a score of association for the ROI. To construct a “null” distribution and assess the strength of any score *τ*\*, we randomly select *N* = 500 other “seed” vertices across the mesh of each *Tarsius* tooth and uniformly generate *N*-“null” regions that are *K*-vertices wide. We then compute similar (null) scores *τ*_1_,…,*τ_N_* for each randomly generated region. A “*P*-value”-like quantity (for the *i*-th molar) is then generated by:

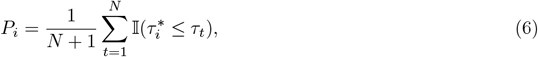

where 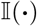 is an indicator function, and a smaller *P_i_* means more confidence in either method’s ability to find the desired paraconid landmark. To ensure the robustness of this analysis, we generate the *N*-random null regions in two ways: *(i)* using a *K*-nearest neighbors (KNN) algorithm on each of the *N*-random seed vertices [56], or *(ii)* manually constructing *K*-vertex wide null regions with surface areas equal to that of the paraconid ROI (Supplementary Material, Section 5). In both settings, we take the median of the *P_i_* values in Equation (6) across all teeth, and report them for each genus and choice of *k* combination — see the first half of Tables 1–2 for the SINATRA implementations and Supplementary Table 3 for results with Limit Shapes, respectively.

**Table 1.**
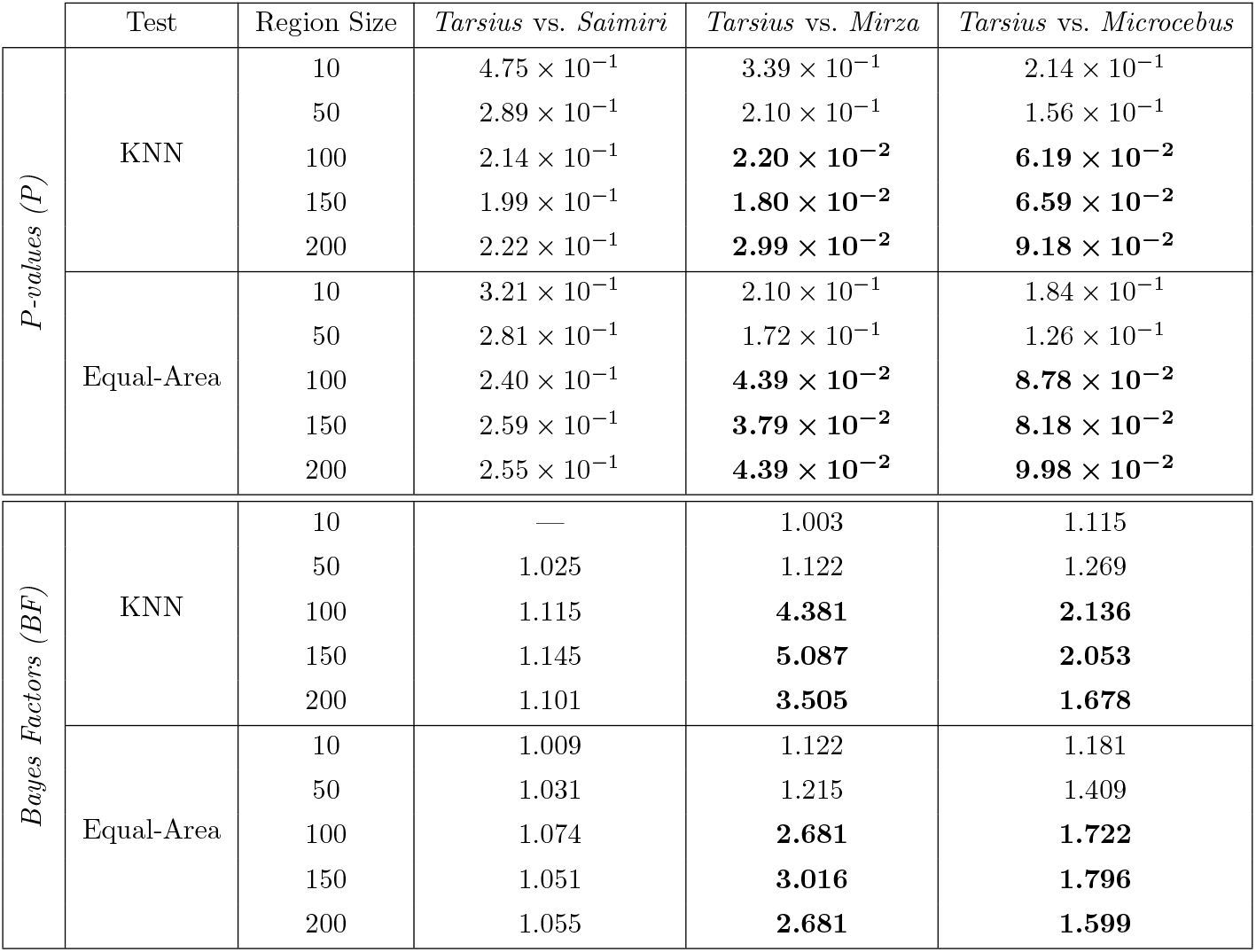
Null region experiment to evaluate SINATRA’s ability to find paraconids in *Tarsius* molars using meshes with correspondence maps. Here, we assess how likely it is that SINATRA finds the region of interest (ROI) by chance. In this experiment, meshes were aligned using the `auto3dgm` software [55]. To produce the results above, we generate 500 “null” regions on each *Tarsius* tooth using *(i)* a KNN algorithm and *(ii)* an equal-area approach (Supplementary Material, Section 5). Next, for each region, we sum the evidence potential or “birth times” of all its vertices. We compare how many times the aggregate scores for the ROI is less than those for the null regions. The median of these “P-values”, and their corresponding calibrated Bayes factors (BF) when median P < 1/e, across all teeth are provided above for the three primate comparisons. Results with values *P*-values less than 0.1 and BFs greater than 1.598 are in bold.

|  | Test | Region Size | *Tarsius* vs. *Saimiri* | *Tarsius* vs. *Mirza* | *Tarsius* vs. *Microcebus* |
| --- | --- | --- | --- | --- | --- |
| *P-values (P)* | KNN | 10 | $4.75 \times 10^{-1}$ | $3.39 \times 10^{-1}$ | $2.14 \times 10^{-1}$ |
| | | 50 | $2.89 \times 10^{-1}$ | $2.10 \times 10^{-1}$ | $1.56 \times 10^{-1}$ |
| | | 100 | $2.14 \times 10^{-1}$ | **$2.20 \times 10^{-2}$** | **$6.19 \times 10^{-2}$** |
| | | 150 | $1.99 \times 10^{-1}$ | **$1.80 \times 10^{-2}$** | **$6.59 \times 10^{-2}$** |
| | | 200 | $2.22 \times 10^{-1}$ | **$2.99 \times 10^{-2}$** | **$9.18 \times 10^{-2}$** |
| | Equal-Area | 10 | $3.21 \times 10^{-1}$ | $2.10 \times 10^{-1}$ | $1.84 \times 10^{-1}$ |
| | | 50 | $2.81 \times 10^{-1}$ | $1.72 \times 10^{-1}$ | $1.26 \times 10^{-1}$ |
| | | 100 | $2.40 \times 10^{-1}$ | **$4.39 \times 10^{-2}$** | **$8.78 \times 10^{-2}$** |
| | | 150 | $2.59 \times 10^{-1}$ | **$3.79 \times 10^{-2}$** | **$8.18 \times 10^{-2}$** |
| | | 200 | $2.55 \times 10^{-1}$ | **$4.39 \times 10^{-2}$** | **$9.98 \times 10^{-2}$** |
| *Bayes Factors (BF)* | KNN | 10 | — | 1.003 | 1.115 |
|  |  | 50 | 1.025 | 1.122 | 1.269 |
|  |  | 100 | 1.115 | **4.381** | **2.136** |
|  |  | 150 | 1.145 | **5.087** | **2.053** |
|  |  | 200 | 1.101 | **3.505** | **1.678** |
|  | Equal-Area | 10 | 1.009 | 1.122 | 1.181 |
|  |  | 50 | 1.031 | 1.215 | 1.409 |
|  |  | 100 | 1.074 | **2.681** | **1.722** |
|  |  | 150 | 1.051 | **3.016** | **1.796** |
|  |  | 200 | 1.055 | **2.681** | **1.599** |

**Table 2.** Null region experiment to evaluate SINATRA’s ability to find paraconids in *Tarsius* molars using meshes without any prior knowledge of correspondence maps. Here, we assess how likely it is that the SINATRA algorithm finds the region of interest (ROI) by chance. In this experiment, we first conduct the Euler characteristic transformation on the raw data for each tooth. Then we *implicitly* normalize the collection of 3D meshes by aligning their EC curves — see the description of approximate grid search and alignment procedure detailed in Section 4 of the Supplementary Material. To produce the results in the table above, we generate 500 “null” regions on each *Tarsius* tooth using *(i)* a KNN algorithm and *(ii)* an equal-area approach (Supplementary Material, Section 5). Next, for each region, we sum the evidence potential or “birth times” of all its vertices. We compare how many times the aggregate scores for the ROI is less than those for the null regions. The median of these “P-values”, and their corresponding calibrated Bayes factors (BF) when median P < 1/e, across all teeth are provided above for the three primate comparisons. Results with values *P*-values less than 0.1 and BFs greater than 1.598 are in bold.

|  | Test | Region Size | *Tarsius* vs. *Saimiri* | *Tarsius* vs. *Mirza* | *Tarsius* vs. *Microcebus* |
| --- | --- | --- | --- | --- | --- |
| *P-values (P)* | KNN | 10 | $3.99 \times 10^{-1}$ | $6.09 \times 10^{-1}$ | $2.06 \times 10^{-1}$ |
| | | 50 | $4.15 \times 10^{-1}$ | $3.23 \times 10^{-1}$ | **$9.58 \times 10^{-2}$** |
| | | 100 | $3.83 \times 10^{-1}$ | $1.72 \times 10^{-1}$ | **$7.59 \times 10^{-2}$** |
| | | 150 | $3.79 \times 10^{-1}$ | $1.16 \times 10^{-1}$ | **$8.38 \times 10^{-2}$** |
| | | 200 | $3.97 \times 10^{-1}$ | $1.52 \times 10^{-1}$ | **$1.00 \times 10^{-1}$** |
| | Equal-Area | 10 | $2.97 \times 10^{-1}$ | $4.49 \times 10^{-1}$ | $1.54 \times 10^{-1}$ |
| | | 50 | $3.37 \times 10^{-1}$ | $3.39 \times 10^{-1}$ | $1.20 \times 10^{-1}$ |
| | | 100 | $3.87 \times 10^{-1}$ | $2.16 \times 10^{-1}$ | **$7.58 \times 10^{-2}$** |
| | | 150 | $4.43 \times 10^{-1}$ | $1.66 \times 10^{-1}$ | **$9.18 \times 10^{-2}$** |
| | | 200 | $4.57 \times 10^{-1}$ | $2.25 \times 10^{-1}$ | **$8.98 \times 10^{-2}$** |
| *Bayes Factors (BF)* | KNN | 10 | — | — | 1.131 |
|  |  | 50 | — | 1.008 | **1.637** |
|  |  | 100 | — | 1.216 | **1.882** |
|  |  | 150 | — | 1.474 | **1.770** |
|  |  | 200 | — | 1.286 | **1.598** |
|  | Equal-Area | 10 | 1.020 | — | 1.278 |
|  |  | 50 | 1.004 | 1.003 | 1.447 |
|  |  | 100 | — | 1.119 | **1.881** |
|  |  | 150 | — | 1.235 | **1.678** |
|  |  | 200 | — | 1.095 | **1.700** |

Using *P*-values as a direct metric of evidence can cause problems. For example, moving from *P* = 0.03 to *P* = 0.01 does not increase evidence for the alternative hypothesis (or against the null hypothesis) by a factor of 3. To this end, we use a calibration formula that transforms a *P*-value to a bound/approximation of a Bayes factor (BF) [21], the ratio of the marginal likelihood under the alternative hypothesis *H*_1_ versus the null hypothesis *H*_0_:

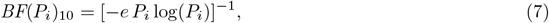

for *P_i_* < 1/*e* and BF(*P_i_*)_10_ is an estimate of 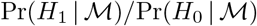, where 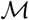 are the molars as meshes and *H*_0_ and *H*_1_ are the null and alternative hypotheses, respectively. The second half of Tables 1–2 and Supplementary Table 3 report these calibrated Bayes factor estimates.

Overall, the Limit Shapes algorithm performs generally well on these data; however, the sparsity in the resulting vertex weights hinders clear visual enrichment of the paraconid in the *Tarsius* molar (e.g., Fig. 7(a)). As a reminder, there is the additional limitation of needing correctly specified point-by-point maps between each tooth in the data to achieve sufficient power. Based on our simulations, if these user inputs had been misspecified, the Limit Shapes approach would have experienced an even more difficult time identifying the region of interest. When using SINATRA on the pre-aligned data, the paraconid ROI is most strongly enriched in the comparisons between the *Tarsius* and either of the strepsirrhine primates, rather than for the *Tarsius-Saimiri* comparison (e.g., Fig. 7(b)). This trend remains consistent even when SINATRA is implemented on the raw data without correspondence maps (e.g., Fig. 7(c)) — although, we note that the enrichment of the ROI is not as distinct which is most likely due to shape alignments being slightly less precise with topological summary statistics versus using known landmarks. We suspect that the difference in enrichment across primate comparisons is partly explained by the divergence times between the genera: *Tarsius* is more recently diverged from *Saimiri* than from the strepsirrhines. This conjecture is consistent with the intuition from our simulation studies, where classes of shapes with sufficiently different morphology result in more accurate identification of unique ROI. On the other hand, the *Tarsius-Saimiri* comparison is analogous to the simulations under to the null model: with too-similar molars, no region appears key to explaining the variance between the two classes of primates.

## Discussion

In this paper, we introduce SINATRA: the first statistical pipeline for sub-image analysis that does not require landmarks or correspondence points between images. We use simulations to demonstrate properties of SINATRA and we illustrate the practical utility of SINATRA on real data. There are many potential extensions to our proposed framework. First, in its current formulation and software implementation, SINATRA is limited to binary classification, but we believe that extensions to multi-class problems and regression with continuous responses are trivial. To analyze continuous traits and phenotypes in many evolutionary applications, one must first disentangle adaptation and heredity [57–60]. The standard approach for this disentanglement is to explicitly account for the hierarchy of descent by adding genetic covariance or kinship across species to the likelihood either via phylogenetic regression [61] or linear mixed models (e.g., the animal model) [62]. Modeling covariance structures also arises in statistical and quantitative genetics applications where individuals are related [63–65]. The SINATRA framework uses a Bayesian hierarchical model that is straightforward to adapt to analyze complex covariance structures in future work.

A second natural extension would be to apply the SINATRA framework to radiomic studies where the goal is to detail the correlation between clinical imaging features and genomic assays [1]. This would require improving the efficacy of SINATRA’s sub-image selection capabilities in data settings where intra-class heterogeneity is high. For example, when studying the progression of glioblastoma multiforme (GBM) — a glioma that materializes into aggressive, cancerous tumor growths within the human brain — magnetic resonance images (MRIs) can vary greatly between patients harboring the same oncogenic mutations. As demonstrated in the current work, SINATRA is generally most powered in scenarios where just a few global features drive the variation between phenotypic classes (e.g., Figs. 2–5 and Supplementary Figs. 1-7). In the examples that we consider here, class definitions are derived from base shapes or governed by some rule of phylogeny. This keeps intra-class heterogeneity low and facilitates the ability of SINATRA to detect varying patterns between shape topologies. As a result, SINATRA is well-suited for anthropologic-based applications and other similar domains. However, in the radiomics context, it is possible that the signature of a given tissue type is made up of small focal lesions that are collectively important in explaining clinical outcomes. Moving forward, more work needs to be done to transfer the utility of SINATRA to cases where inter-class variation is driven by local fluctuations in shape morphology.

## Code Availability

Code for implementing the SINATRA pipeline is freely available at https://github.com/lcrawlab/SINATRA, and is written in `R` (version 3.5.3). As part of this procedure: *(i)* inference for the Gaussian process classification (GPC) model using elliptical slice sampling was carried out using the `R` package `FastGP` (version 1.2) [66] and *(ii)* the computation of effect sizes and association measures for the Euler characteristic curves was done with the “RelATive cEntrality (RATE)” source code in `R` (version 1.0.0; https://github.com/lorinanthony/RATE) [39]. Visualizing the reconstructed regions outputted by SINATRA was done using the package `rgl` (version 0.100.19) [67], and general utility functions for triangular meshes from the package `Rvcg` (version 0.18) [68]. Furthermore, preprocessing steps for the meshes examined in the study were performed using `Morpho` (version 2.60) [68,69] and `auto3dgm` (Version 1.00) [55].

## Data Availiability

The current study makes use of two real shape datasets. The first consists of Lemuridae teeth, a specific genera of Cercopithecidae (Old World monkeys; http://www.wisdom.weizmann.ac.il/~ylipman/CPsurfcomp/) [11]. The second is comprised of mandibular molars from two different suborders of the primate: Haplorhini (“dry-nosed” primates; https://gaotingran.com/codes/codes.html) and Strepsirrhini (“moist-nosed” primates; http://morphosource.org/Detail/ProjectDetail/Show/project_id/89). From the first suborder, we have 33 molars from the *Tarsius* [11,50,51] and 9 molars from the *Saimiri* [52] genera. In the second suborder, we have 11 molars from the *Microcebus* and 6 molars from the *Mirza* genera [11, 50, 51]. In the initial analysis, the meshes of all teeth were aligned using `auto3dgm` [55]. This algorithm establishes correspondences between uniformly placed landmarks on each tooth such that each mesh has the same orientation (e.g., Supplementary Fig. 8). After alignment, the molars were translated to be centered at the origin and normalized to be enclosed within a unit ball. In a secondary analyses, we demonstrate how to implement SINATRA in settings where *a priori* correspondences between shapes are unavailable. Here, we first conduct the Euler characteristic transformation on the raw data for each tooth and then we “implicitly” normalize the meshes by aligning the EC curves (e.g., Supplementary Figs. 9 and 10). Details on the approximate grid search procedure used to do this is given in full detail in Section 4 of the Supplementary Material. These `auto3dgm` and ECT quality controlled meshes were then used to demonstrate the utility of the SINATRA pipeline, respectively.

## Acknowledgements

We would like to thank the Editor, Associate Editor and two anonymous referees for their constructive comments. We would also like to thank Ani Eloyan, Anthea Monod, Jenny Tung, Katharine Turner, and Christine Wall for helpful conversations and suggestions. We are also very appreciative to Ruqi Huang and Maks Ovsjanikov for sharing their `MATLAB` implementation of Limit Shapes. This research was partly supported by grants P20GM109035 (COBRE Center for Computational Biology of Human Disease; PI Rand) and P20GM103645 (COBRE Center for Central Nervous; PI Sanes) from the NIH NIGMS, 2U10CA180794-06 from the NIH NCI and the Dana Farber Cancer Institute (PIs Gray and Gatsonis), as well as by an Alfred P. Sloan Research Fellowship. A majority of this research was conducted using computational resources and services at the Center for Computation and Visualization (CCV), Brown University. SM would like to acknowledge partial funding from HFSP RGP005, NSF DMS 17-13012, NSF BCS 1552848, NSF DBI 1661386, NSF IIS 15-46331, NSF DMS 16-13261, as well as high-performance computing partially supported by grant 2016-IDG-1013 from the North Carolina Biotechnology Center. Lastly, TG was supported by NSF Grant No. DMS-1439786 while in residence at the Institute for Computational and Experimental Research in Mathematics (ICERM) in Providence, RI, during the Computer Vision Semester Program in Spring 2019, and partially by NSF DMS-1854831 afterwards. Any opinions, findings, and conclusions or recommendations expressed in this material are those of the author(s) and do not necessarily reflect the views of any of the funders.

## Author Contributions Statement

LC conceived the study. SM and LC developed the methods. BW, TS, and HK developed the algorithms and implemented the software. DB designed sampling strategy for the molar analysis. TG constructed the shape caricaturization and provided the baseline of correspondence-based methods. All authors performed the analyses, interpreted the results, and wrote and revised the manuscript.

