## Supplementary Text, Figures, and Tables for "A Statistical Pipeline for Identifying Physical Features that Differentiate Classes of 3D Shapes"

### 1 Technical Results

#### 1.1 Posterior Inference via Elliptical Slice Sampling

This section details how we conduct posterior inference for the Gaussian process classification (GPC) model [1–4]. Recall our goal is to take observations from two different shape classes and identify physical characteristics that best detail the variation between them. In the main text, we assume an $(n \times p)$-dimensional design matrix $\mathbf{X}$ of topological summary statistics with the following hierarchical model

$$\mathbf{y} \sim \mathcal{B}(\boldsymbol{\pi}), \qquad \mathrm{g}(\boldsymbol{\pi}) = \Phi^{-1}(\boldsymbol{\pi}) = \boldsymbol{f}, \qquad \boldsymbol{f} \sim \mathcal{N}(\mathbf{0}, \mathbf{K}) \tag{1}$$

where $\mathbf{y}$ is an $n$-dimensional vector of Bernoulli independently distributed class labels, $\boldsymbol{\pi}$ is an $n$-dimensional vector representing the underlying probability that a shape is classified as a "case" (i.e., $y_i = 1$), $\mathrm{g}(\cdot)$ is a probit link function with $\Phi(\cdot)$ the cumulative distribution function (CDF) of the standard normal distribution, and $\boldsymbol{f}$ is an $n$-dimensional vector which needs to be estimated.

In this specification, we assume $\boldsymbol{f}$ follows a multivariate normal distribution with mean vector $\mathbf{0}$ and a covariance matrix $\mathbf{K}$ computed using a Gaussian kernel function. Given the complete specification of the GPC, our goal is to draw samples from the posterior distribution of the latent variables $\boldsymbol{f}$. Using Bayes' theorem, begin by considering

$$p(\boldsymbol{f} \,|\, \mathbf{y}) \propto p(\mathbf{y} \,|\, \boldsymbol{f}) p(\boldsymbol{f}), \tag{2}$$

where $p(\mathbf{y} \,|\, \boldsymbol{f})$ denotes the likelihood of the observed binary labels given the functions (i.e., the Bernoulli distribution), and $p(\boldsymbol{f})$ is the prior distribution for the latent variables (i.e., the multivariate normal distribution).

The probit likelihood in Equation (1) makes it intractable to estimate the posterior distribution $p(\boldsymbol{f} \,|\, \mathbf{y})$ via closed-form solution. We instead use a Markov chain Monte Carlo (MCMC) method called "elliptical slice sampling" to conduct posterior inference [5]. This technique is similar to a Metropolis-Hastings algorithm [6] in that samples from a target distribution are found via some "transition" or "proposal" kernel. In this work, we consider the parametrization

$$\boldsymbol{f}^* = \boldsymbol{f} \cos\vartheta + \mathbf{z} \sin\vartheta, \tag{3}$$

where the auxiliary variable $\mathbf{z}$ defines a full ellipse passing through a current state $\boldsymbol{f}$, parameterized by a step size $\vartheta$. The elliptical slice sampling algorithm works by selecting a new state $\boldsymbol{f}^*$ (on the ellipse) using the transition kernel described in Equation (3). Unlike Metropolis-Hastings, there is no formal rejection of $\boldsymbol{f}^*$ since the new state equals the current state $\boldsymbol{f}$ only if it is the only location on the ellipse with nonzero likelihood. In the event of an unacceptable $\boldsymbol{f}^*$, the algorithm proposes a new step size $\vartheta \in [\vartheta_{\min}, \vartheta_{\max}]$, where the bounds are iteratively (and adaptively) shrunk until an acceptable new state $\boldsymbol{f}^*$ is found. An overview of the elliptical slice sampling scheme is detailed in Algorithm 1 below.

#### 1.2 Closed Form Solution for Relative Centrality Association Measures

This section details how we compute evidence of association measures for each topological summary statistic. After implementing the elliptical slice sample algorithm to draw samples from the posterior distribution of the latent variables $\boldsymbol{f}$, we define a nonparametric effect size for each topological summary statistic via the following linear transformation [7–9]

$$\boldsymbol{\beta} = (\mathbf{X}^{\intercal}\mathbf{X})^{-1}\mathbf{X}^{\intercal}\boldsymbol{f}, \tag{4}$$

where each element in $\boldsymbol{\beta}$ details the relationship between the Euler characteristic features and the variance between shape classes. We use these effect sizes to assign an entropic-based measure of relative centrality

**Algorithm 1** Elliptical Slice Sampling for Gaussian Process Classification

**Input:** A current state sampled from $\boldsymbol{f} \sim \mathcal{N}(\mathbf{0}, \mathbf{K})$, and a log-likelihood function $\log p(\mathbf{y} \mid \boldsymbol{f})$.
**Output:** A new state $\boldsymbol{f}^*$ with a marginal distribution proportional to the target posterior distribution.

```
Generate an ellipse by sampling auxiliary variables: z ~ N(0, K).
Draw b ~ U[0, 1] from a standard uniform distribution and define a log-likelihood threshold:
        T ← log p(y | f) + log b.
Draw an initial proposal ϑ ~ U[0, 2π] and define an interval:
        [ϑ_min, ϑ_max] ← [ϑ − 2π, ϑ].
Generate a new state: f* = f cos ϑ + z sin ϑ.
if log p(y | f) > T then:
    Accept the new state f*.
else:
    Shrink the step size interval, and sample a new point:
    if ϑ < 0 then: ϑ_min ← ϑ.
    else: ϑ_max ← ϑ.
    end if
    ϑ ~ U[ϑ_min, ϑ_max].
    GoTo 4.
end if
```

to each $j$-th topological feature using Kullback-Leibler divergence (KLD) [10–17]

$$\mathrm{KLD}(\beta_j) := \mathrm{KL}\left[p(\boldsymbol{\beta}_{-j}) \,\|\, p(\boldsymbol{\beta}_{-j} \mid \beta_j = 0)\right] = \int_{\boldsymbol{\beta}_{-j}} \log\left(\frac{p(\boldsymbol{\beta}_{-j})}{p(\boldsymbol{\beta}_{-j} \mid \beta_j = 0)}\right) p(\boldsymbol{\beta}_{-j})\, \mathrm{d}\boldsymbol{\beta}_{-j}. \tag{5}$$

As mentioned in the main text, we study the difference between *(i)* the conditional posterior distribution $p(\boldsymbol{\beta}_{-j} \mid \beta_j = 0)$ with the effect of the $j$-th topological feature being set to zero, and *(ii)* the marginal posterior distribution $p(\boldsymbol{\beta}_{-j})$ with the effects of the $j$-th feature being integrated out. The KLD is non-negative, and equals zero if and only if the $j$-th topological feature is of little importance, since removing its effect has no influence on the other features. For simplicity, we assume that the implied posterior distribution of $\boldsymbol{\beta}$ (deterministically given in Equation (4)) is approximately multivariate normal with an empirical mean vector $\boldsymbol{\mu}$ and positive semi-definite covariance/precision matrix $\boldsymbol{\Sigma} = \boldsymbol{\Lambda}^{-1}$. Given these values, we iteratively partition such that, for each $j$-th topological feature:

$$\boldsymbol{\beta} = \begin{pmatrix} \beta_j \\ \boldsymbol{\beta}_{-j} \end{pmatrix}; \quad \boldsymbol{\mu} = \begin{pmatrix} \mu_j \\ \boldsymbol{\mu}_{-j} \end{pmatrix}; \quad \boldsymbol{\Sigma} = \begin{pmatrix} \sigma_j & \boldsymbol{\sigma}_{-j}^{\intercal} \\ \boldsymbol{\sigma}_{-j} & \boldsymbol{\Sigma}_{-j} \end{pmatrix}; \quad \boldsymbol{\Lambda} = \begin{pmatrix} \lambda_j & \boldsymbol{\lambda}_{-j}^{\intercal} \\ \boldsymbol{\lambda}_{-j} & \boldsymbol{\Lambda}_{-j} \end{pmatrix}. \tag{6}$$

Under normality assumptions, Equation (5) has the following closed form solution

$$\mathrm{KLD}(\beta_j) = \frac{1}{2}\left[-\log|\boldsymbol{\Sigma}_{-j}\boldsymbol{\Lambda}_{-j}| + \mathrm{tr}(\boldsymbol{\Sigma}_{-j}\boldsymbol{\Lambda}_{-j}) + 1 - p + \alpha_j(\beta_j - \mu_j)^2\right], \tag{7}$$

where $\log|\cdot|$ represents the matrix log-determinant function, and $\mathrm{tr}(\cdot)$ is the matrix trace function. Importantly, the term $\alpha_j = \boldsymbol{\lambda}_{-j}^{\intercal}\boldsymbol{\Lambda}_{-j}^{-1}\boldsymbol{\lambda}_{-j}$ characterizes the linear (and non-negative) rate of change of information when the effect of any topological feature is absent from the analysis [9]. By symmetry in the notation for elements of the sub-vectors and sub-matrices, we simply permute the order of the variables in $\boldsymbol{\beta}$ and iteratively compute the KLD to measure the centrality of each Euler characteristic transform. Finally, we normalize by $\gamma_j = \text{KLD}(\beta_j)/\sum \text{KLD}(\beta_l)$, as this is bounded on the unit interval $[0,1]$ with the natural interpretation of providing relative evidence of association for shape features.

In practice, we use a few approximations to scale the otherwise computationally expensive steps in Equation (7). The first approximation involves computing the log determinant. With a dataset of reasonably dense 3D shapes, the number of topological features is expected to be large (i.e., $p \gg 0$). In this setting, the term $-\log(|\boldsymbol{\Sigma}_{-j}\boldsymbol{\Lambda}_{-j}|) + \text{tr}(\boldsymbol{\Sigma}_{-j}\boldsymbol{\Lambda}_{-j}) + (1-p)$ remains relatively equal for each feature $j$ and makes a negligible contribution to the entire sum. Thus, we simplify Equation (7) to

$$\text{KLD}(\widetilde{\beta}_j) \approx \alpha_j(\widetilde{\beta}_j - \mu_j)^2/2. \tag{8}$$

This approximation of the KLD still relies on the full precision matrix $\boldsymbol{\Lambda}$. For large number of topological features $p$, this calculation is expensive; however, it is only done once and is used for all $p$ computations. The rate of change parameter $\alpha_j = \boldsymbol{\lambda}_{-j}^{\intercal}\boldsymbol{\Lambda}_{-j}^{-1}\boldsymbol{\lambda}_{-j}$, on the other hand, depends on the partitioned matrix $\boldsymbol{\Lambda}_{-j}^{-1}$ for every $j$-th topological feature. This requires inverting a $(p-1)\times(p-1)$ matrix $p$ times. Fortunately, we can reduce this computational burden by taking advantage of the fact that any $\boldsymbol{\Lambda}_{-j}^{-1}$ is formed by removing the $j$-th row and column from the precision matrix $\boldsymbol{\Lambda}$. Therefore, given the partition in Equation (6), we can use the Sherman-Morrison formula [18] to efficiently approximate these quantities using the following rank-1 update for each topological feature

$$\boldsymbol{\Omega}^{(j)} = \boldsymbol{\Lambda} - \boldsymbol{\Lambda}\boldsymbol{\sigma}_j\boldsymbol{\sigma}_j^{\intercal}\boldsymbol{\Lambda}/\left(1+\boldsymbol{\sigma}_j^{\intercal}\boldsymbol{\Lambda}\boldsymbol{\sigma}_j\right) \qquad j=1,\ldots,p. \tag{9}$$

Here, $\boldsymbol{\sigma}_j$ is the $j$-th column from the posterior covariance matrix $\boldsymbol{\Sigma}$, and each $\boldsymbol{\Lambda}_{-j}^{-1}$ is approximated by removing the $j$-th row and column from $\boldsymbol{\Omega}^{(j)}$. Ultimately, this reduces the computational complexity of Equation (8) to just $p$-independent $O(p^2)$ operations which can be parallelized.

#### 1.3 Reconstruction: Identifying Vertices via Critical Point Equations

In this section, we detail how we identify vertices from corresponding topological summary statistics during shape reconstruction. Let $\{\boldsymbol{\nu}_1,\ldots,\boldsymbol{\nu}_d\}$ with $\boldsymbol{\nu}_i = (\nu_{i,1},\nu_{i,2},\nu_{i,3})$ be the $d$-directions that map onto the same $m$-set of unknown vertices on a shape $\{\mathbf{q}_1,\ldots,\mathbf{q}_m\}$ with $\mathbf{q}_t = (x_t,y_t,z_t)$. By definition, each Euler characteristic curve is calculated along a given direction with a predefined filtration that changes at $m$-heights $\{\mathbf{h}_1,\ldots,\mathbf{h}_m\}$ with $\mathbf{h}_t = (h_{t,1},\ldots,h_{t,d})$, accounting for possible multiplicities [19, 20]. This results in the system of equations

$$\begin{bmatrix} \nu_{1,1} & \nu_{1,2} & \nu_{1,3} \\ \nu_{2,1} & \nu_{2,2} & \nu_{2,3} \\ \vdots & \vdots & \vdots \\ \nu_{d,1} & \nu_{d,2} & \nu_{d,3} \end{bmatrix} \begin{bmatrix} x_1 & x_2 & \cdots & x_m \\ y_1 & y_2 & \cdots & y_m \\ z_1 & z_2 & \cdots & z_m \end{bmatrix} = \begin{bmatrix} h_{1,1} & h_{1,2} & \cdots & h_{1,m} \\ h_{2,1} & h_{2,2} & \cdots & h_{2,m} \\ \vdots & \vdots & \ddots & \vdots \\ h_{d,1} & h_{d,2} & \cdots & h_{d,m} \end{bmatrix}, \tag{10}$$

which may be interpreted as an equation representing $d$-planes that intersect at $m$-points in $\mathbb{R}^3$. We want to recover the unknown vertices from known pairs of directions and height functions $(\boldsymbol{\nu},\mathbf{h})$. In practice, Equation (10) may not have a unique solution depending on the number of directions $d$. For example, when $d=3$, there are $(m!)^2$ solutions that correspond to permuting the second and third direction with respect to the first. As long as $m$ is bounded, there exists a finite $d$ that guarantees a unique solution: the minimal number of directions such that collection of $d$-parallel planes intersect at exactly $m$-points [21]. This leads to the following proposition.

**Proposition 1.** *Let $\{\boldsymbol{\nu}_1,\ldots,\boldsymbol{\nu}_d\}$ be a collection of $d$ distinct directions contained within a cone of small enough cap radius $\theta$ that detect the same set of $m$-critical points of a mesh. Also assume that the mesh* *contains no collection of three collinear vertices. If $d = d(m)$ is sufficiently large, then d-fold intersections of the hyperplanes determined by $\{\boldsymbol{\nu}_1, \ldots, \boldsymbol{\nu}_d\}$ are the coordinates of the critical points detected by each $\boldsymbol{\nu}_i$.*

*Proof.* We first prove that if $d = 3$, then there are at most $(m!)^2$ point configurations that satisfy the critical point equation, with equality if and only if all directions detect critical vertices at distinct heights. The configurations are given by first matching the heights of $\boldsymbol{\nu}_1$ to $\boldsymbol{\nu}_2$, which can be done in $m!$ ways. We can match the $m$-heights of $\boldsymbol{\nu}_3$ to these matched pairs in $m!$ ways, yielding $(m!)^2$ possible configurations.

Second, we show that if $d$ directions admit $k > 1$ configurations, then there exists $D$ such that $(d+D)$ directions admit fewer than $k$ configurations. Let $\mathcal{S}_1, \ldots, \mathcal{S}_n$ be a set of Euler characteristic curves that detect $m$-critical vertices and define $\mathcal{T}$ to be the set of all vertices in all point configurations admissible by $\{\mathcal{S}_i\}_{i=1}^n$. Let $\mathcal{Q} \subset \mathcal{T}$ be some set of vertices in $\mathcal{T}$ with undetermined coordinates and cardinality $|\mathcal{Q}| = r > 0$.

Now, consider an additional set of Euler characteristic curves $\mathcal{S}_{n+1}, \ldots, \mathcal{S}_{r(r-1)/2+1}$. Since the mesh does not contain a collection of three collinear vertices, any hyperplane associated with any curve can pass through at most two vertices in $\mathcal{Q}$. Since there are only $\binom{r}{2}$ such pairs in $\mathcal{Q}$ and the directions $\{\boldsymbol{\nu}_1, \ldots, \boldsymbol{\nu}_d\}$ are non-antipodal, then by the pigeonhole principle there is some $\mathcal{S}^*$ and vertex $\mathbf{q} \in \mathcal{Q}$ such that a hyperplane associated with $\mathcal{S}^*$ does not pass through $\mathbf{q}$. This means $\mathbf{q}$ is not a critical vertex. Hence, the admissible point configuration of $\mathcal{S}_1, \ldots, \mathcal{S}_n$ that contains $\mathbf{q}$ is an inadmissible configuration for $\mathcal{S}_{n+1}, \ldots, \mathcal{S}_{r(r-1)/2+1}$. □

In the SINATRA pipeline, we work with discretized Euler characteristic curves such that topological features are represented as direction-height bands $(\boldsymbol{\nu}, \mathbf{h})$ of the form

$$\mathcal{S}_{\nu,h} = \{\mathbf{q} \in \mathbb{R}^3 \,|\, \mathbf{h} - \Delta\mathbf{h} \leq \boldsymbol{\nu} \times \mathbf{q} \leq \mathbf{h} + \Delta\mathbf{h}\}, \tag{11}$$

where $\Delta$ details the width of the band. To do shape reconstruction, we combine information within bands from Equation (11) using an idea similar to the theoretical critical point equation in Equation (10). The problem with the critical point equation is that it is not computationally feasible to maintain very fine height discretization and infer exact 3D coordinates. Finer discretization leads to an increased number of observed topological features and, thus requires more directions of analysis to have a sufficiently fine epsilon or $\epsilon$-net to detect each vertex.

To this end, we derive an approximate critical point equation that is scalable enough to be used in practice. From previous work, a generic set of directions detects the same set of vertices when they are sufficiently close to each other: a consequence of *stratification* (see Proposition 6.1 of [19]). For our approximate critical point equation, we use a coarser stratification than previously introduced in the literature. By limiting our scope to groups of directions near each other, we implicitly obtain a set of directions with information that can be combined for statistical inference and visualization on the 3D shape (see Definition 7.3 and Theorem 7.1 of [21]). Now let $\{\boldsymbol{\nu}_1, \ldots, \boldsymbol{\nu}_d \,|\, \theta\}$ denote a cone of $d$ directions with "significant" topological features tabulated by heights $\{\mathbf{h}_1, \ldots, \mathbf{h}_m\}$, where $\theta$ represents the cap radius from which the directions are generated. The reconstructed areas (or vertices) associated with the cone are given by

$$\mathcal{R} := \bigcap_{i=1}^{d} \bigcup_{t=1}^{m} \mathcal{S}_{\nu_i,h_t}. \tag{12}$$

To summarize as much variation across shape classes as possible, we need to run Euler characteristic filtrations along sufficiently many directions — that is, we need to take enough cones to cover the surface of the sphere [19, 21]. As previously suggested, information corresponding to areas with high curvature is easy to observe.

##### 1.4 Brief Note: Intuition Behind using Cones

We describe how SINATRA chooses sets of directions during the computation of Euler characteristic curves. Denote this set of directions as the union of $c$ cones $\mathcal{D} = \bigcup_r \mathcal{C}_r(\theta)$ with $r = 1, \ldots, c$. For consistency, define a cone as $\mathcal{C}_r(\theta) = \{\nu_{r,1}, \ldots, \nu_{r,d} \,|\, \theta\}$, parameterized by an angle $\theta$ from which equidistant vectors are generated from a central direction.

In the SINATRA algorithm, different cones are centered around some predetermined number of equidistributed central directions on the sphere. We use cones because local shape information matters most when determining reconstructed manifolds [21–23]. As previously noted, Euler characteristic curves measured in directions of close proximity contain similar local information. This similarity naturally leads to the construction of sets $\mathcal{C}_r(\theta)$, where the angle $\theta$ between them is small. Ideally, one would select an effectively large number of cones (and directions within these cones) to ensure that SINATRA is covering all shapes in the study and summarizing all relevant information about the variance between classes.

However, there is a direct tradeoff between the number of cones placed on a shape ($c$), and the number of topological summary statistics used to represent its features ($p$). Remember, the dimensionality of the design matrix $\mathbf{X}$ is $n \times p$ with $p = c \times d \times l$, where $d$ is the number of directions within each cone and $l$ is the number of sublevel sets (i.e., filtration steps) to compute the Euler characteristic curve along a given direction. Hence, the problem can become very high-dimensional as $c$ increases.

#### 2 Performance Assessment for Simulations

In the main text, we demonstrate the power of our pipeline for association mapping-based tasks via multiple simulations studies using a sequential procedure:

1. Fit the GPC model using elliptical slice sampling and compute relative centrality association measures $\gamma_j$ for each $j$-th topological feature (i.e., Euler characteristic per sublevel set filtration). Recall, the total number of features $p = c \times d \times l$ is a product of *(i)* $c$, the number of cones of directions; *(ii)* $d$, the number of directions within each cone; and *(iii)* $l$, the number of sublevel sets (i.e., steps in the filtration) used to compute the Euler characteristic (EC) along a given direction.

2. Sort the topological features from largest to smallest according to their association measures $\gamma_1 \geq \gamma_2 \geq \cdots \geq \gamma_p$.

3. By iteratively moving through the sorted measures $T_k = \gamma_k$ (starting with $k = 1$), we reconstruct the vertices corresponding to the topological features with $\{j : \gamma_j \geq T_k\}$.

A vertex is "detected" when the sublevel set in which it resides is selected across all the directions within a particular cone. We form a union of the set of detected vertices across all cones to construct the set of reconstructed vertices at a given level $T_k$, as given in Eq. (12). Using this set of vertices, we compute the true positive rate (TPR) and false positive rate (FPR) by assessing overlap with the set of truly associated vertices used to generate the two classes of simulated shapes:

$$\text{TPR} = \frac{\sum \text{TP}}{\sum \text{P}}, \qquad \text{FPR} = \frac{\sum \text{FP}}{\sum \text{N}} \tag{13}$$

where TP is the number of correctly detected true vertices, P is the total number of causal vertices, TN stands for the true negatives detected by our pipeline, and N stands for the total number of non-causal vertices. In this manner, we obtain a receiver operating characteristic (ROC) curve for the simulation studies. For the synthetic study with perturbed spheres, true vertices correspond to the red cusps or indentations unique to classes 1 and 2, respectively (e.g., Figs. 2(a)-2(c) in the main text). For the caricatured teeth, true vertices correspond to the red exaggerated regions unique to classes 1 and 2, respectively (e.g., Fig. 3(a) in the main text).

### 3 Details Behind Caricature Simulations

In the main text, we carry out a simulation study where we modify triangular mesh representations of Lemuridae teeth using a popular caricaturization procedure [24]. Here, we select a single tooth and create two shape classes (25 shapes per classes and 50 shapes in total) by smoothly modifying pre-specified landmarks to create class-defining regions. These caricatured teeth are then analyzed by SINATRA and we assess its ability to identify the truly associated vertices.

First, we compute a planar parametrization of the 3D mesh using a discrete conformal map [25]. Since each triangular face is in one-to-one correspondence with a planar triangle, we can compute a unique affine transformation from each planar triangle to each triangular face. The collection of these transformations can be viewed as the gradient field of the piecewise linear embedding of the planar region into the ambient, three-dimensional Euclidean space. Such gradient fields are in one-to-one correspondences with the piecewise linear embeddings [26, 27].

We obtain caricatured teeth by simply perturbing the gradient field — that is, we multiply each affine transformation by a positive scalar and then solve a Poisson equation to recover (up to a global rigid motion) the $(x, y, z)$-coordinates of all the vertices in the planar parametrization. Details about setting up and solving the Poisson equations can be found in [24]. Code for implementing these simulations can be found at `https://github.com/lcrawlab/SINATRA`.

### 4 Running Sinatra without Correspondence Maps

In this section, we detail how to implement the SINATRA pipeline without relying on *a priori* correspondence maps between shapes. Here, shapes are commonly considered to be subsets of $\mathbb{R}^3$ modulo rigid motions (i.e., compositions of translations, scaling, rotations and reflections). One way to mod out these motions is via landmarks. This is achieved by first centering the meshes at the origin and scaling them to have the same unit area. Then one proceeds by choosing $k$-landmarks, and seeks to find the rotations and reflections that minimize the distance between sets of landmarks on each mesh. Landmarks are unnecessary for SINATRA because the Euler characteristic transformation is $O(d)$-equivariant:

**Theorem 4.1** (Theorem 6.6 of Turner et al. (2014) [19])**.** *Let $K_1$ and $K_2$ be generic geometric simplicial complexes in $\mathbb{R}^d$. Let $\mu$ be Lesbesgue measure on the of the unit circle or sphere $S^{d-1}$. If $ECT(K_1)_*(\mu) = ECT(K_1)_*(\mu)$ (that is the pushforward of the measures are the same), then there is some $\phi \in O(d)$ such that $K_2 = \phi(K1)$. In other words, that is to say that $K_2$ is some combination of rotations and reflections of $K_1$.*

The key to our results in this section stems directly from the proof of this theorem which tells us that the ECT is an $O(3)$ equivariant transformation. More specifically, for an element $\phi$ in the symmetry group $O(3)$, the following diagram commutes:

$$\begin{array}{ccc} \mathcal{M}_1 & \xrightarrow{\phi} & \mathcal{M}_2 \\ \downarrow{\scriptstyle ECT} & & \downarrow{\scriptstyle ECT} \\ \chi_\nu(\mathcal{M}_1) & \xrightarrow{\phi} & \chi_\nu(\mathcal{M}_2) \end{array}$$

where the EC curve $\chi_\nu(\mathcal{M})$ tracks the change in the Euler characteristic with respect to a given filtration of length $l$ in direction $\nu$. The practical implication from this diagram is that we need not to find any correspondences between the shapes themselves. Instead, after translating and scaling the meshes, we can align the data simply using the Euler curves.

#### 4.1 How to Align Euler Characteristic Curves

Let $\mathcal{M}_1, \ldots, \mathcal{M}_n$ represent the $n$ meshes in our study and denote $\boldsymbol{\nu} = \{\nu_1, \ldots, \nu_D\}$ to be a set of $D$ directions on $S^2$. We will use $\mathrm{ECT}(\mathcal{M}_r)$ to represent the Euler characteristic transformation of the $r$-th mesh $\mathcal{M}_r$ restricted to the finite set of directions $\{\nu_1, \ldots, \nu_D\}$, and we will use $\phi_{r,s}$ for the $O(3)$ action that minimizes the distance between $\mathrm{ECT}(\mathcal{M}_r)$ and $\mathrm{ECT}(\mathcal{M}_s)$. Unlike landmarks, Euler characteristic curves cannot be rotated without knowing the full transformation. However, if the set of directions $\boldsymbol{\nu}$ form an orbit under the subgroup generated by $\phi$, then any rotation in this group corresponds to a permutation $p : [D] \to [D]$. Unfortunately, this is in general not the case, and the rotation is often unknown *a priori*. Instead, we carefully choose the directions $\boldsymbol{\nu}$ such that we can approximate any rotation by a permutation of the Euler characteristic curves with some error.

To accomplish this goal, note that the element $\phi$ can be represented as a combination of rotations along the $x$- and $y$- and $z$-axes, with respective angles $\theta, \delta, \psi \in [0, 2\pi]$ and the corresponding reflection $x \mapsto -x$. To recover $\phi$, we perform a grid search over $O(3)$. First, we let $g$ denote the size of the grid for each of the axis-aligned rotations. Then, for $\mathrm{sign}_l \in \{-1, 1\}$ and $\theta_i, \delta_j, \psi_k \in [0, 2\pi/n, \ldots, 2\pi]$, we write

$$A_{ijkl} = \begin{pmatrix} \mathrm{sign}_l & 0 & 0 \\ 0 & \mathrm{sign}_l & 0 \\ 0 & 0 & \mathrm{sign}_l \end{pmatrix} \begin{pmatrix} \cos(\psi_k) & -\sin(\psi_k) & 0 \\ \sin(\psi_k) & \cos(\psi_k) & 0 \\ 0 & 0 & 1 \end{pmatrix} \begin{pmatrix} \cos(\theta_i) & 0 & \sin(\theta_i) \\ 0 & 1 & 0 \\ -\sin(\theta_i) & 0 & \cos(\theta_i) \end{pmatrix} \begin{pmatrix} 1 & 0 & 0 \\ 0 & \cos(\delta_j) & -\sin(\delta_j) \\ 0 & \sin(\delta_j) & \cos(\delta_j) \end{pmatrix}$$

Here, each rotation $A_{ijkl}$ is associated with a permutation $p_{ijkl} : [D] \to [D]$ on the scanning directions. The associated permutation is computed as the solution to the linear assignment problem

$$p_{ijkl} = \operatorname*{arg\,min}_{p:[D]\to[D]} \sum_{t=1}^{D} d(\nu_{p(t)} - A_{ijkl}\nu_t), \tag{14}$$

where the distance $d$ between any two directions $\nu_1$ and $\nu_2$ is given by the squared Euclidean distance

$$d(\nu_1, \nu_2) = \|\nu_1 - \nu_2\|_2^2. \tag{15}$$

The alignment of Euler characteristic curves reduces to finding the transformation $\widetilde{A}_{12}$ that corresponds to the permutation $\widetilde{p}$ that satisfies

$$\widetilde{p} = \operatorname*{arg\,min}_{p\in\mathcal{P}} d(\mathrm{ECT}(\mathcal{M}_1), p \times \mathrm{ECT}(\mathcal{M}_2)) \tag{16}$$

where $\mathcal{P} = \bigcup p_{ijkl} \subset S_D$ is the set of permutations of $[D]$ that are minimizers for the rotation problem in Equation (14).

#### 4.2 Proof-of-Concept Alignment Case Study

We study the performance of post transformation alignment procedure in a case where the ground truth is known. Namely, we took a random subset of five mandibular molars from the primate dataset highlighted in the main text, and compared the alignments done with the Euler characteristic curves against the state-of-the-art quasi Branch and Bound shape alignment method [28], to which we input 300 landmarks per mesh that were picked by `auto3dgm` using the hybrid furthest point and Gaussian process sampling. We take the EC curves along $d = 2918$ directions, $l = 200$ sublevel sets per direction, and conduct the grid search along 16,000 elements of $O(3)$. The alignment results for this toy dataset are presented in Figure 9 where there is great agreement between the two approaches. Figure 10 shows the meshes of all teeth in the primate dataset using the aligned Euler curves.

### 5 Generating Null Regions in *Tarsius* Paraconid Analysis

In the main text, we perform an analysis to detect distinct morphological features between different genera of primates, with *Tarsius* molars the class of interest. Morphologically, tarsier teeth are seemingly tritubercular and have an additional cusp, allowing them to eat a wider range of foods reasonably well [29]. To this end, we perform three pairwise comparisons with each of the other genera (*Tarsius* versus *Saimiri*, *Mirza*, and *Microcebus*, respectively), and assess how likely it is that SINATRA and the Limit Shapes algorithm [30] finds the paraconid region of interest (ROI) by chance.

To ensure the robustness of this analysis, we generate the $N$-random null regions in two ways: *(i)* using a $K$-nearest neighbors (KNN) algorithm on each of the $N$-random seed vertices [31], or *(ii)* manually constructing the null regions with some $K$-nearest vertices such that they have surface areas approximately equal to that of the paraconid ROI. In the latter setting, we ensure the appropriate surface area for the enclosed null region around a given seed vertex by using the following procedure:

(1) Propose different $K$-vertex wide neighborhoods until we find some integer $K_0$ and $(K_0+1)$ such that the surface area of the ROI lies somewhere between them.

(2) Identify the "circumferential" vertices which lie inside the $(K_0+1)$-neighborhood but outside the $K_0$-neighborhood of the seed vertex, and compute the surface area assigned to each of these vertices.

(3) Sort the surface area values assigned to the circumferential vertices in descending order. Let $N$ be the smallest integer such that the surface area of the region formed by adding the first $N$-circumferential vertices in the ordering exceeds the area of the ROI.

The desired null region is marked by the union of the vertices in the $K_0$-neighborhood and the $N$-circumferential vertices in step (3).

### Supplementary Figures

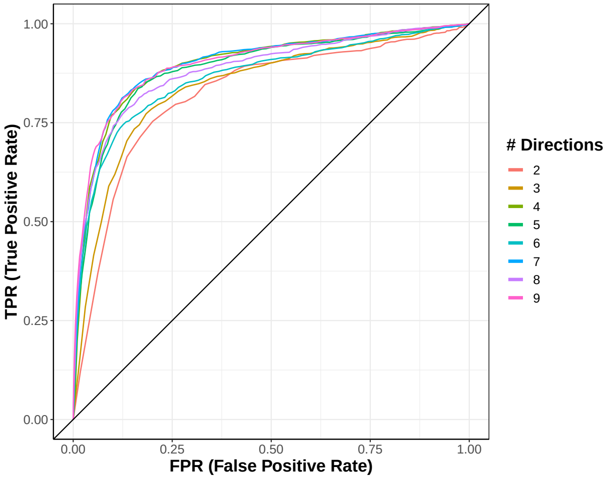

(a) Scenario I

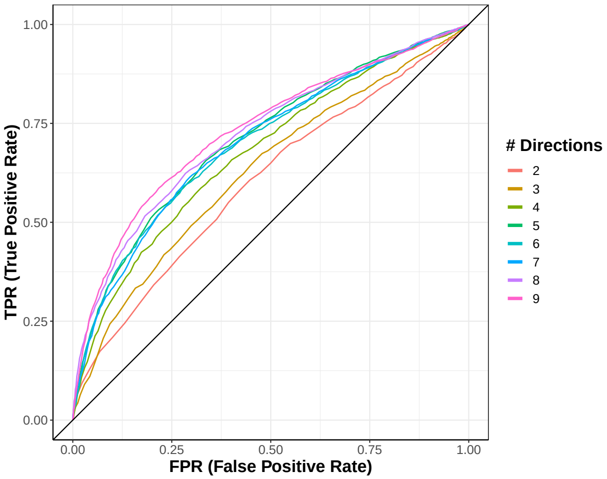

(b) Scenario II

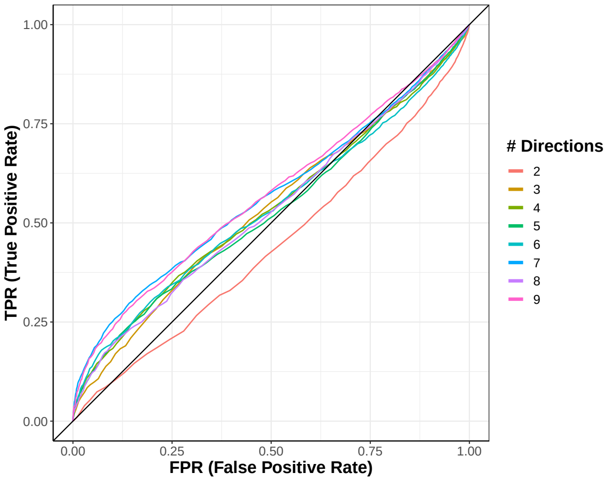

(c) Scenario III

**Supplementary Figure 1. Power and sensitivity analysis for detecting associated regions while changing the number of directions $d$ taken within each cone.** For this analysis, we generate 100 shapes by partitioning unit spheres into regions 10 vertices-wide, centered at 50 equidistributed points. Two shape classes (50 shapes per class) are defined by shared and class-specific characteristics. The shared or "non-associated" features are chosen by randomly selecting $u$ regions and pushing the sphere outward at each of these positions. This is done for all shapes, regardless of class. To generate class-specific or "associated" features, $v$ distinct regions are chosen for a given class and perturbed inward. We vary these parameters and analyze three increasingly more difficult simulation scenarios: **(a)** $u = 2$ shared and $v = 1$ associated; **(b)** $u = 6$ shared and $v = 3$ associated; and **(c)** $u = 10$ shared and $v = 5$ associated. In these panels, ROC curves depict the overall ability of SINATRA to identify vertices located within these associated regions, as a function of increasing the number of directions taken within a given cone. The main takeaway from this sensitivity analysis is consistent with both intuition and theoretical results from the literature [8, 19, 21]: seeing more of a shape (i.e., using more directions) generally leads to an improved ability to map back onto associated regions. Other SINATRA parameters were fixed: $c = 25$ cones, $\theta = 0.15$ cap radius used to generate directions in a cone, and $l = 30$ sublevel sets per filtration. Results are based on 50 simulated replicates.

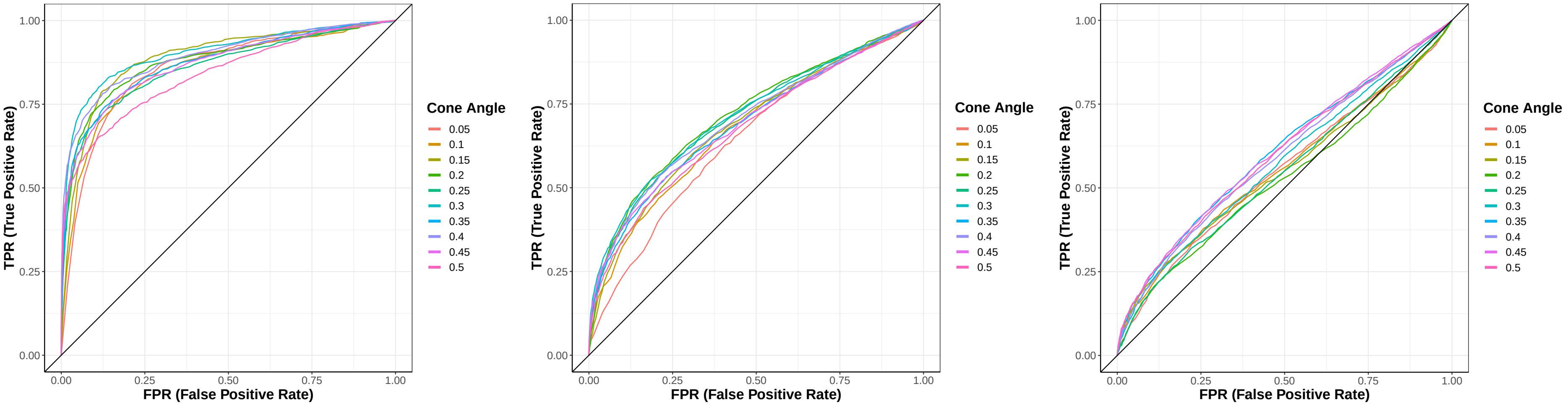

(a) Scenario I (b) Scenario II (c) Scenario III

**Supplementary Figure 2. Power and sensitivity analysis for detecting associated regions while changing the angle $\theta$ between directions within cones.** For this analysis, we generate 100 shapes by partitioning unit spheres into regions 10 vertices wide, centered at 50 equidistributed points. Two shape classes (50 shapes per class) are defined by shared and class-specific characteristics. The shared or "non-associated" features are chosen by randomly selecting $u$ regions and pushing the sphere outward at each of these positions. To generate class-specific or "associated" features, $v$ distinct regions are chosen for a given class and perturbed inward. We vary these parameters and analyze three increasingly more difficult simulation scenarios: **(a)** $u = 2$ shared and $v = 1$ associated; **(b)** $u = 6$ shared and $v = 3$ associated; and **(c)** $u = 10$ shared and $v = 5$ associated. In these panels, ROC curves depict the overall ability of SINATRA to identify vertices located within these associated regions, as a function of increasing "generative cone angle" or the distance between individual directions within a given cone. The main takeaway from this sensitivity analysis is that cones should be defined by directions in close proximity to each other; but not so close that they effectively contain the exact same local information. Other SINATRA parameters were fixed: $c = 25$ cones, $d = 5$ directions per cone, and $l = 30$ sublevel sets per filtration. Results are based on 50 simulated replicates.

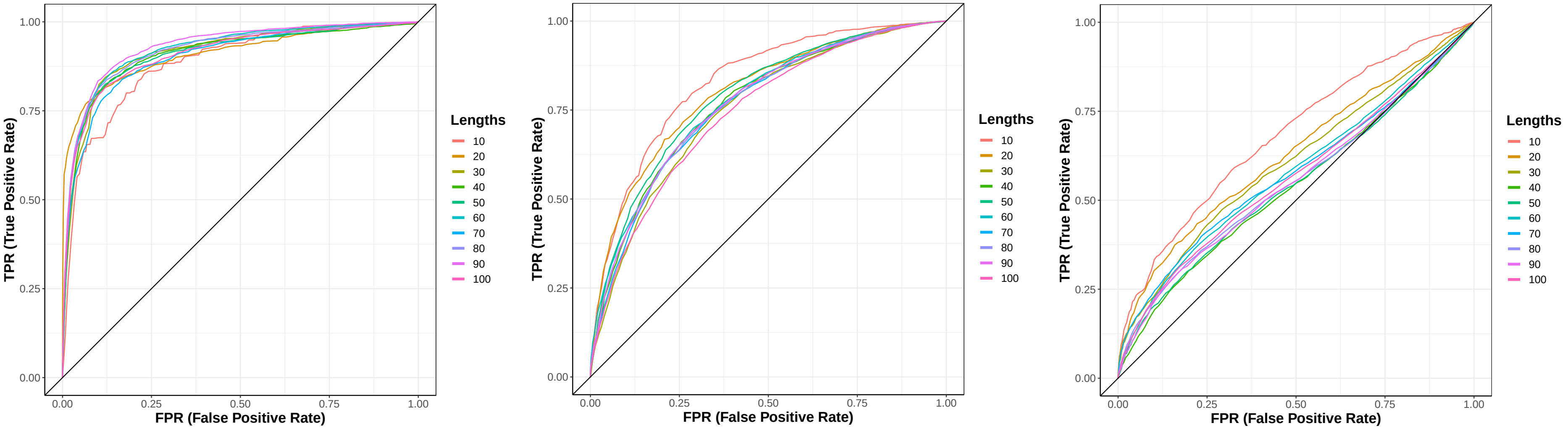

(a) Scenario I (b) Scenario II (c) Scenario III

**Supplementary Figure 3. Power and sensitivity analysis for detecting associated regions while changing the number sublevel sets $l$ per filtration.** For this analysis, we generate 100 shapes by partitioning unit spheres into regions 10 vertices wide, centered at 50 equidistributed points. Two shape classes (50 shapes per class) are defined by shared and class-specific characteristics. The shared or "non-associated" features are chosen by randomly selecting $u$ regions and pushing the sphere outward at each of these positions. To generate class-specific or "associated" features, $v$ distinct regions are chosen for a given class and perturbed inward. We vary these parameters and analyze three increasingly more difficult simulation scenarios: **(a)** $u = 2$ shared and $v = 1$ associated; **(b)** $u = 6$ shared and $v = 3$ associated; and **(c)** $u = 10$ shared and $v = 5$ associated. In these panels, ROC curves depict the overall ability of SINATRA to identify vertices located within these associated regions, as a function of increasing the granularity of the steps during the filtration over the shapes. For intricate shapes, choosing improperly scaled steps during the computation of Euler characteristic curves can cause the algorithm to miss relevant information and lose power. If only a few features define shapes, this loss is less important. Other SINATRA parameters were fixed: $c = 25$ cones, $d = 5$ directions per cone, and $\theta = 0.15$ cap radius used to generate directions in a cone. Results are based on 50 simulated replicates.

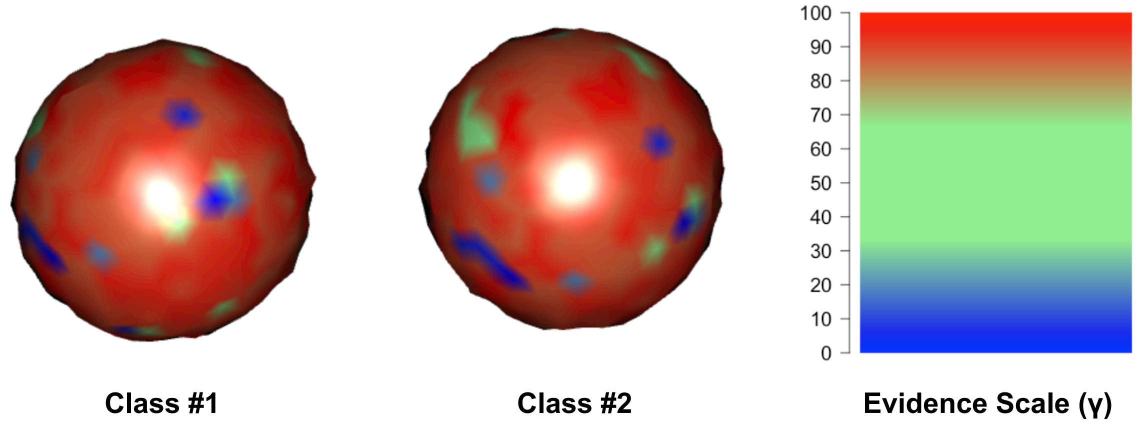

(a) Two Classes of Effectively Matching Shapes

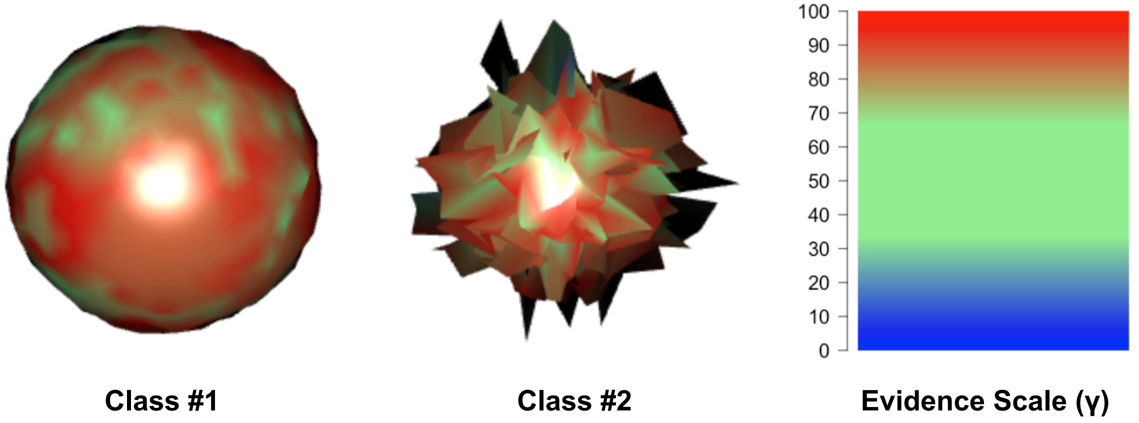

(b) Two Classes of Completely Dissimilar Shapes

**Supplementary Figure 4. Simulation results with shapes generated under the null hypothesis of relative centrality.** These proof-of-concept simulations showcase the two scenarios where SINATRA fails to identify meaningful reconstructed regions: **(a)** two classes of shapes that are effectively the same, and **(b)** two classes of shapes that are completely dissimilar. In both scenarios, all features contribute equally to the variation between the classes. In **(a)**, shapes in both classes appear to be same (up to some small Gaussian noise); therefore, there are no "associated" regions and no group of vertices distinctively stand out. In **(b)**, shapes between the two classes look nothing alike. In this case, all vertices are seemingly associated and no one feature is central or key to explaining the observed variation. The colors display vertex evidence potential on a scale from $[0 - 100]$. A maximum of 100 represents the threshold at which the first shape vertex is reconstructed, while 0 denotes the threshold when the last vertex is reconstructed. Regardless of the scenario, since no meaningful regions differentiate between two classes of shapes, (mostly) all vertices appear to be born relatively early and at the same time. Results are based on taking 50 classes per shape class.

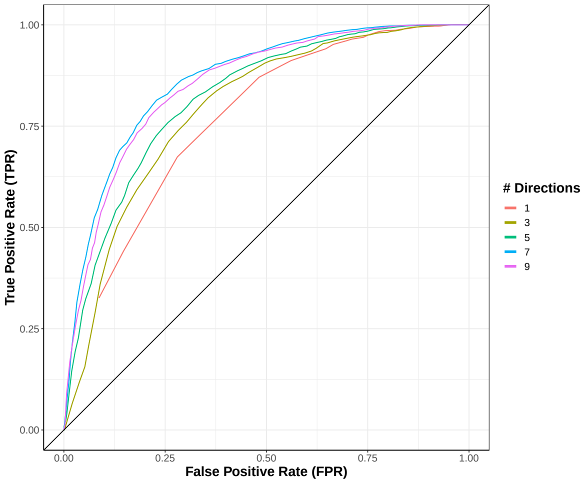

(a) Scenario I

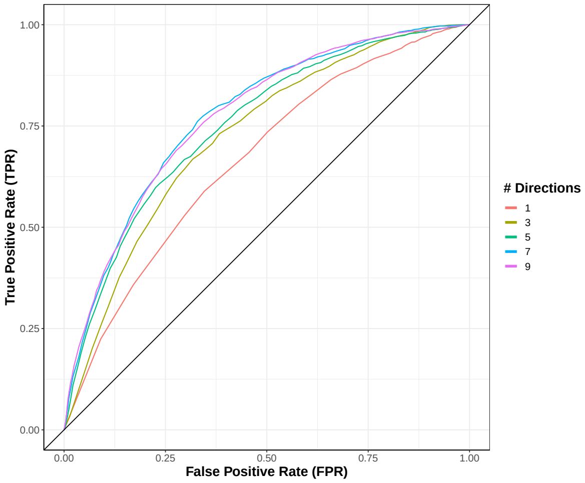

(b) Scenario II

**Supplementary Figure 5. Power and sensitivity analysis for detecting associated vertices across different classes of caricatured shapes while changing the number of directions $d$ taken within each cone.** We modify real Lemuridae molars using a common caricaturization procedure. First, we fix the triangular mesh of an individual tooth. Next, we take known landmarks for the tooth [32], and assign $v$ of them to one class and $v'$ to the other. The caricaturization multiplies each face within these regions by positive scalars so that class-specific features are exaggerated. We repeat twenty-five times (with some small added noise) to create two classes of 25 shapes. SINATRA analyzes the synthetic shapes to identify the associated regions. We consider two scenarios by varying the number of class-specific landmarks that determine the caricaturization. In scenario I, we set $v, v' = 3$; and in scenario II, $v, v' = 5$. In panels **(a)** and **(b)**, ROC curves depict the ability of SINATRA to identify vertices located within associated regions, as a function of the number of directions taken within a given cone. Seeing more of a shape (i.e., using more directions) generally leads to an improved ability to map back onto associated regions. Other SINATRA parameters were fixed: $c = 15$ cones, $\theta = 0.15$ cap radius used to generate directions in a cone, and $l = 50$ sublevel sets per filtration. Results are based on 50 replicates in each scenario.

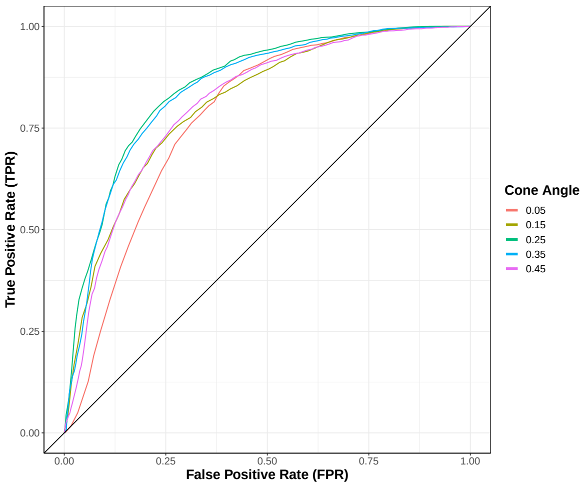

(a) Scenario I

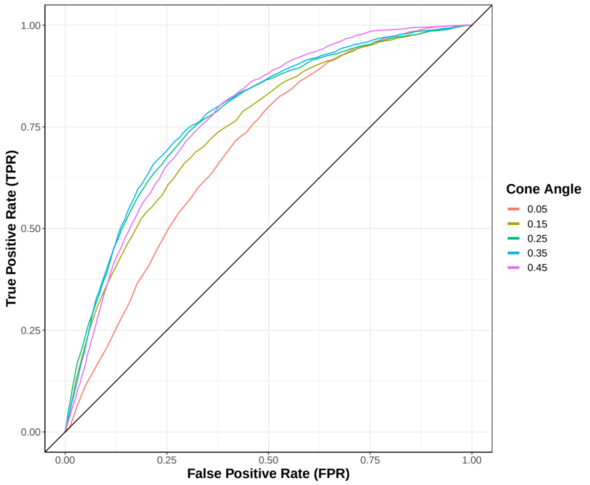

(b) Scenario II

**Supplementary Figure 6. Power and sensitivity analysis for detecting associated vertices across different classes of caricatured shapes while changing the angle $\theta$ between directions within cones.** We modify real Lemuridae molars using a widely used caricaturization procedure. First, we fix the triangular mesh of an individual tooth. Next, we take known landmarks for the tooth [32], and assign $v$ of them to one class and $v'$ to the other. The caricaturization multiplies each face within these regions by positive scalars to exaggerate class-specific features. We repeat twenty-five times (with some small added noise) to create two classes of 25 shapes. SINATRA analyzes the synthetic shapes to identify the associated regions. We consider two scenarios by varying the number of class-specific landmarks that determine the caricaturization. In scenario I, we set $v, v' = 3$; and in scenario II, $v, v' = 5$. In panels **(a)** and **(b)**, ROC curves depict the ability of SINATRA to identify vertices located within associated regions, as a function of "generative cone angle" or the distance between individual directions within a given cone. Cones should be defined by directions that are in some optimal proximity to each other. Other SINATRA parameters were fixed: $c = 15$ cones, $d = 5$ directions per cone, and $l = 50$ sublevel sets per filtration. Results are based on 50 replicates in each scenario.

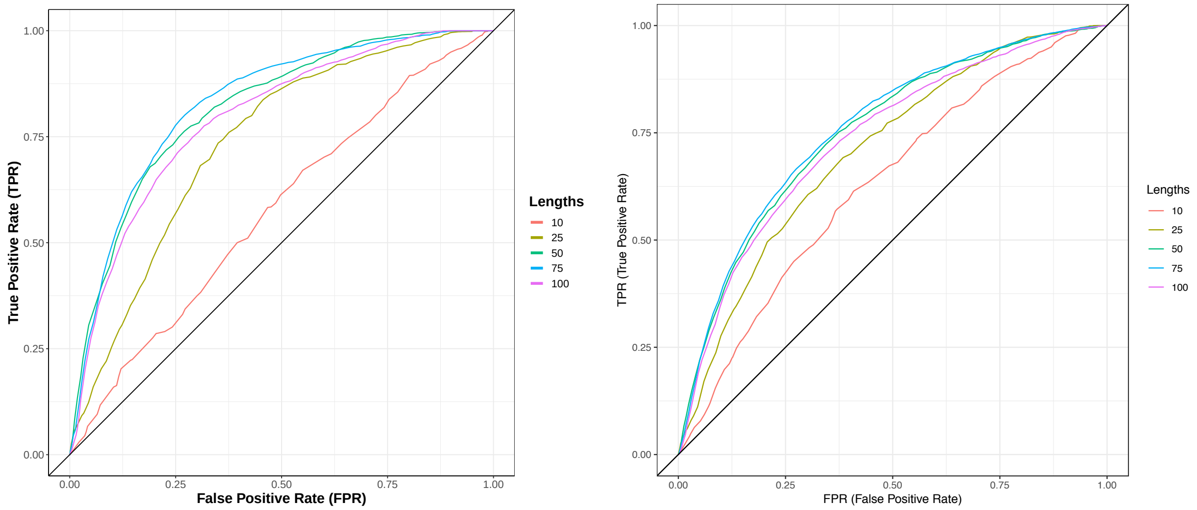

(a) Scenario I

(b) Scenario II

**Supplementary Figure 7. Power and sensitivity analysis for detecting associated vertices across different classes of caricatured shapes while changing the number sublevel sets $l$ per filtration.** We modify real Lemuridae molars using a common caricaturization procedure. First, we fix the triangular mesh of an individual tooth. Next, we take known landmarks for the tooth [32], and assign $v$ of them to one class and $v'$ to the other. The caricaturization multiplies each face within these regions by positive scalars to exaggerate class-specific features. We repeat twenty-five times (with some small added noise) to create two classes of 25 shapes. SINATRA analyzes the synthetic shapes to identify the associated regions. We consider two scenarios by varying the number of class-specific landmarks that determine the caricaturization. In scenario I, we set $v, v' = 3$; and in scenario II, $v, v' = 5$. In panels **(a)** and **(b)**, ROC curves depict the ability of SINATRA to identify vertices located within associated regions, as a function of the granularity of the steps during the filtration over the shapes. For intricate shapes, choosing an improper filtration step size can cause the algorithm to miss important information and lose power. Other SINATRA parameters were fixed: $c = 15$ cones, $d = 5$ directions per cone, and and $\theta = 0.15$ cap radius used to generate directions in a cone. Results are based on 50 replicates in each scenario.

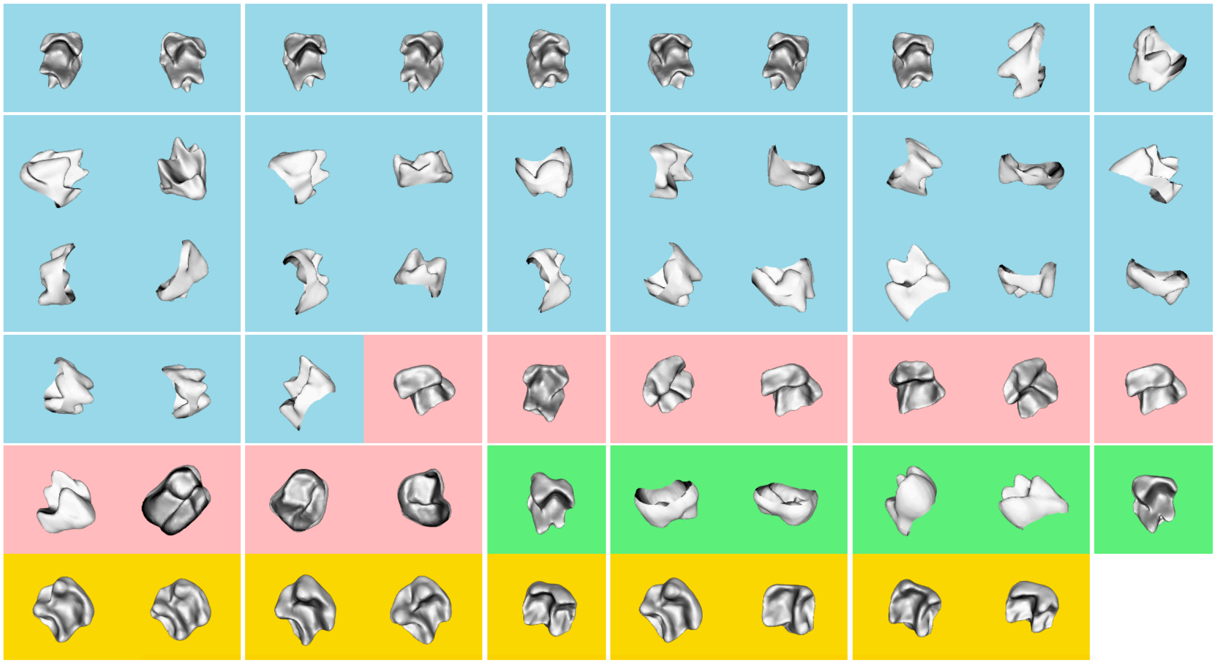

(a) Unaligned Version of Teeth (Raw Data)

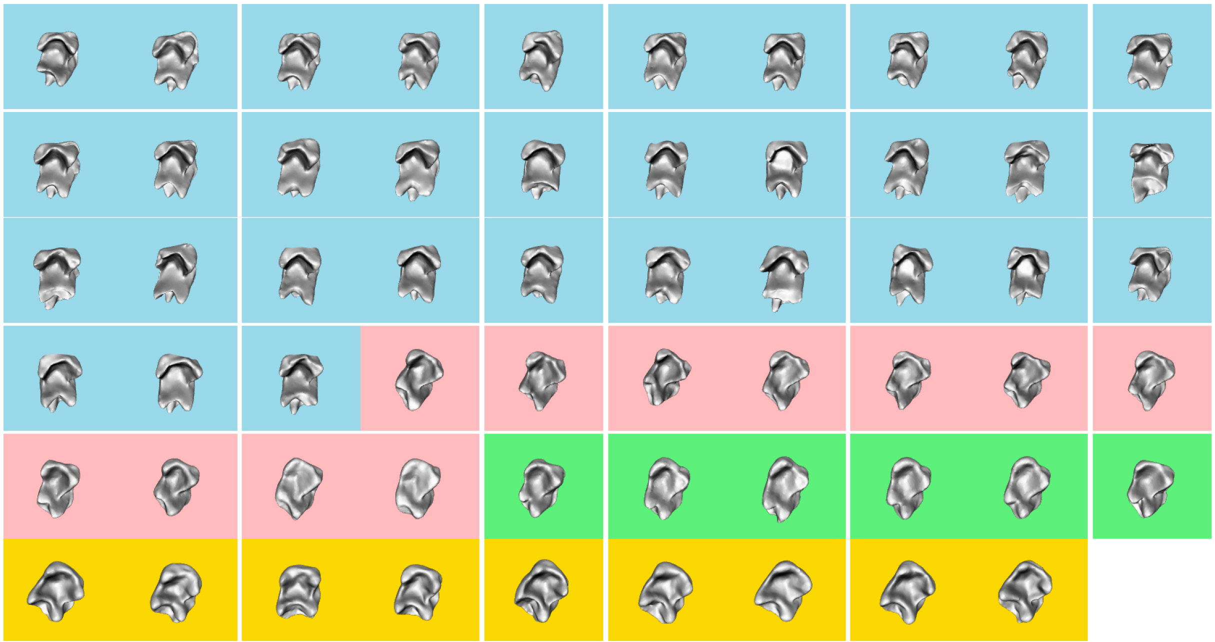

(b) `Autodgm` Aligned Version of Teeth (Quality Controlled Data)

**Supplementary Figure 8. Visualization of 3D data before and after quality control and alignment procedures.** We display mandibular molars from two different suborders of the primate: Haplorhini ("dry-nosed" primates) and Strepsirrhini ("moist-nosed" primates). In the first suborder, 33 molars come from the *Tarsius* (blue) and and 9 molars from the *Saimiri* (gold) genera. In the second suborder, 11 molars come from the *Microcebus* (pink) and 6 molars from the *Mirza* (green) genera. Panel **(a)** depicts the raw data, while **(b)** shows the meshes of all teeth post-alignment using the `auto3dgm` software [33].

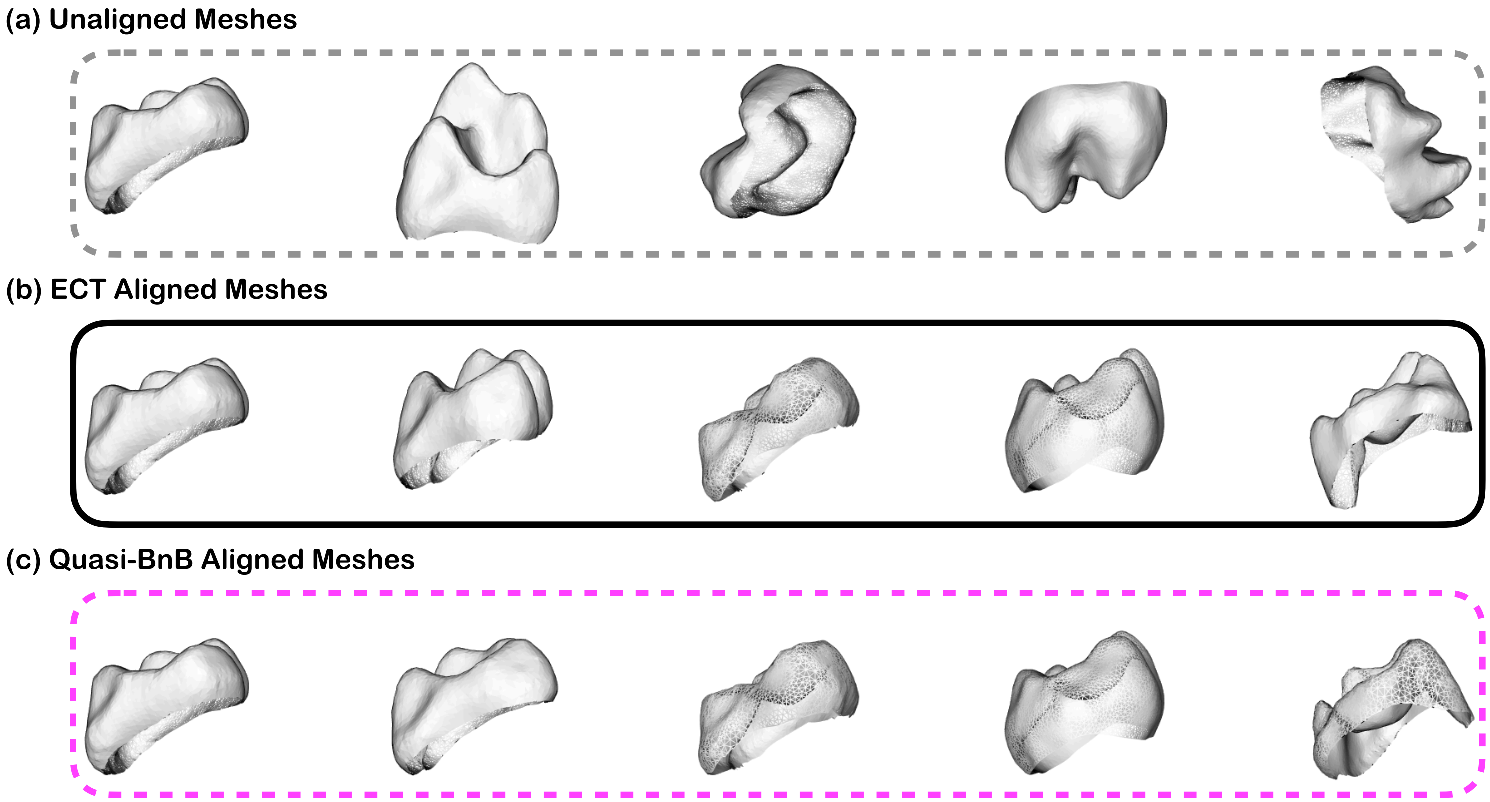

**Supplementary Figure 9. Results from the proof-of-concept study demonstrating the performance of aligning meshes by aligning Euler characteristic curves.** Here, we randomly select a random subset of five mandibular molars taken from the primate dataset analyzed in the main text. Panel **(a)** depicts the raw data. Panel **(b)** shows meshes that have been aligned without the use of correspondences between shapes. In this procedure, we first conduct the Euler characteristic transformation on the raw data for each tooth. Then we implicitly normalize the meshes by aligning the EC curves via the approximate grid search procedure detailed in Section 4 of the Supplementary Material. To generate this figure, we take the EC curves along $d = 2918$ directions, $l = 200$ sublevel sets per direction, and conduct the grid search along 16,000 elements of $O(3)$. As baseline comparison, panel **(c)** shows the performance of the state-of-the-art quasi Branch and Bound (BnB) shape alignment method [28], to which we input 300 landmarks per mesh that were picked by `auto3dgm` using the hybrid furthest point and Gaussian process sampling. Notably, we see significant agreement between the two approaches.

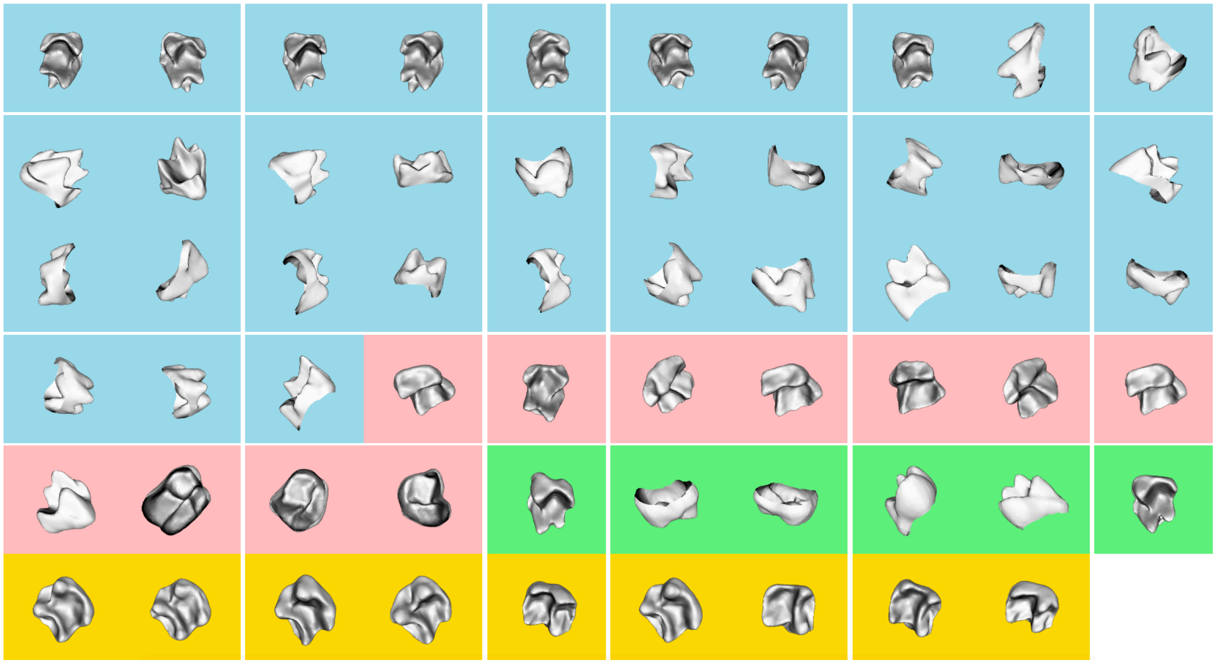

(a) Unaligned Version of Teeth (Raw Data)

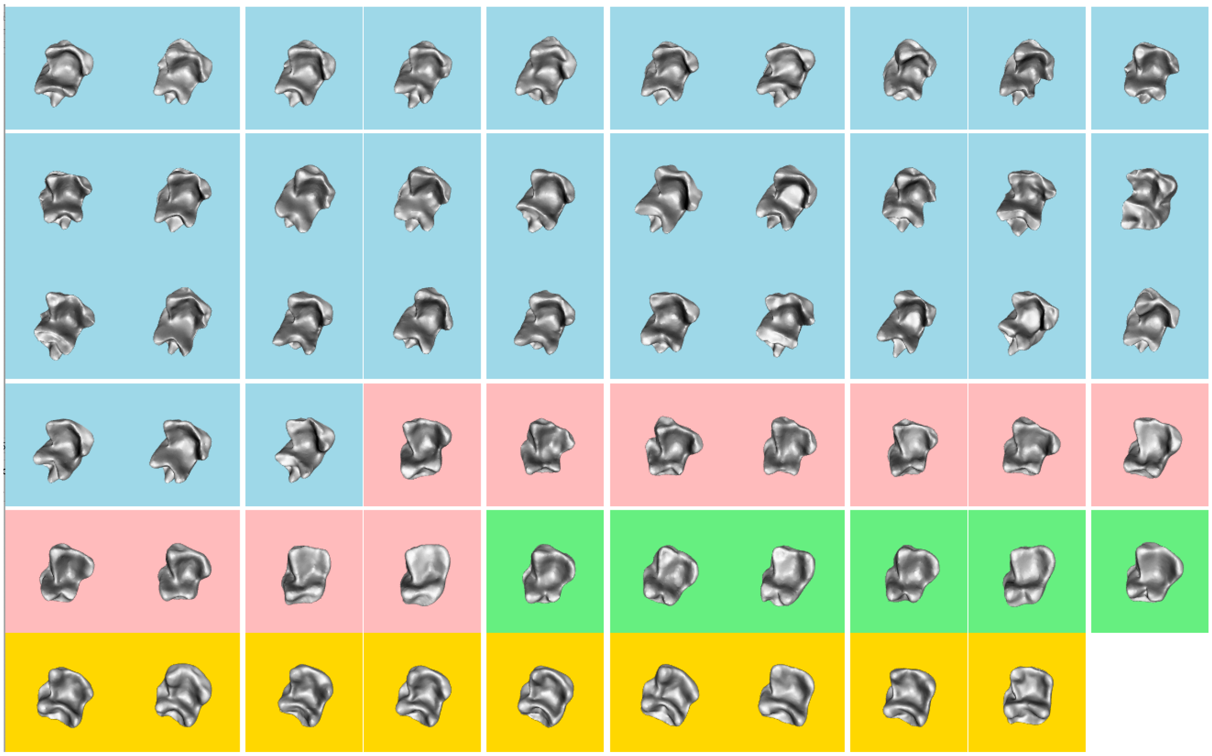

(b) ECT Aligned Version of Teeth (Quality Controlled Data without Correspondences)

**Supplementary Figure 10. Visualization of 3D data before and after quality control and alignment procedures.** We display mandibular molars from two different suborders of the primate: Haplorhini ("dry-nosed" primates) and Strepsirrhini ("moist-nosed" primates). In the first suborder, 33 molars come from the *Tarsius* (blue) and and 9 molars from the *Saimiri* (gold) genera. In the second suborder, 11 molars come from the *Microcebus* (pink) and 6 molars from the *Mirza* (green) genera. Panel **(a)** depicts the raw data, while **(b)** shows the meshes of all teeth using the aligned Euler curves as detailed in Section 4 of the Supplementary Material.

### Supplementary Tables

| Free Parameters in SINATRA Software | | | |
|---|---|---|---|
| **Notation** | **Description** | **Range** | **General Guidelines** |
| $r$ | Radius of the bounding sphere for the shapes | — | Usually set to 1/2 since meshes are assumed to be scaled to the unit sphere |
| $c$ | Number of cones of directions | $[1, \infty)$ | Set greater than 1 as more power can be achieved by taking filtrations over multiple directions |
| $d$ | Number of directions per cone | $[1, \infty)$ | Set greater than 1 as more power can be achieved by taking filtrations over multiple directions |
| $\theta$ | Cap radius used to generate directions within a cone | $(0, 2\pi]$ | Set between $[0.15, 0.25]$ since cones should be defined by directions in close proximity |
| $l$ | Number of sublevel sets (filtration steps) | $[1, \infty)$ | Optimal choice depends on the types of shapes being analyzed so use grid search |

**Supplementary Table 1. General guidelines for choosing values for the free parameters in the SINATRA pipeline software.** The SINATRA algorithm requires the following inputs: *(i)* aligned shapes represented as meshes; *(ii)* $\mathbf{y}$, a binary vector denoting shape classes; *(iii)* $r$, the radius of the bounding sphere for the shapes; *(iv)* $c$, the number of cones of directions; *(v)* $d$, the number of directions within each cone; *(vi)* $\theta$, the cap radius used to generate directions in a cone; and *(vii)* $l$, the number of sublevel sets (i.e., filtration steps) to compute the Euler characteristic (EC) along a given direction. The guidelines provided are based off of intuitions gained through the simulation studies provided in the main text. In practice, we suggest specifying multiple cones $c > 1$ and utilizing multiple directions $d$ per cone (see monotonically increasing power in Figs. 2-3 in the main text and Supplementary Figs. 1-7). While the other two parameters ($\theta$ and $l$) do not have monotonic properties, their effects on SINATRA's performance still have natural interpretations. Selection of $\theta \in [0.15, 0.25]$ supports previous theoretical results that cones should be defined by directions in close proximity to each other [21]; but not so close that they explain the same local information with little variation. As we show in the next section, optimal choice of $l$ depends on the types of shapes being analyzed. Intuitively, for very intricate shapes, coarse filtrations with too few sublevel sets cause the algorithm to miss or "step over" very local undulations in a shape. In practice, we recommend choosing the angle between directions within cones $\theta$ and the number of sublevel sets $l$ via cross validation or some grid-based search.

| **Average Run Time (sec)** | | | | | |
|---|---|---|---|---|---|
| | | | Number of Shapes ($n$) | | |
| # of Cones ($c$) | Directions per Cone ($d$) | Sublevel Sets ($l$) | $n = 25$ | $n = 50$ | $n = 100$ |
| $c = 25$ | $d = 5$ | $l = 25$ | 61.07 (1.48) | 120.97 (0.72) | 196.14 (1.41) |
| | | $l = 50$ | 106.25 (4.27) | 251.80 (1.92) | 438.04 (1.75) |
| | $d = 10$ | $l = 25$ | 138.92 (2.11) | 312.30 (1.53) | 528.24 (1.67) |
| | | $l = 50$ | 294.19 (1.73) | 773.01 (6.14) | 1521.87 (3.18) |
| $c = 50$ | $d = 5$ | $l = 25$ | 137.93 (2.43) | 306.34 (1.52) | 533.66 (1.95) |
| | | $l = 50$ | 294.15 (3.21) | 783.17 (7.06) | 1549.92 (7.54) |
| | $d = 10$ | $l = 25$ | 354.88 (3.54) | 890.51 (9.49) | 1668.77 (5.79) |
| | | $l = 50$ | 965.47 (9.75) | 2894.31 (34.68) | 6683.44 (33.72) |
| $c = 75$ | $d = 5$ | $l = 25$ | 237.93 (5.05) | 560.77 (3.54) | 1017.33 (3.52) |
| | | $l = 50$ | 580.36 (8.81) | 1639.21 (16.05) | 3449.41 (9.77) |
| | $d = 10$ | $l = 25$ | 683.05 (8.72) | 1831.22 (14.29) | 3675.09 (17.91) |
| | | $l = 50$ | — | — | — |

**Supplementary Table 2. Empirical times for running the SINATRA algorithm as a function of its free parameters and inputs.** Each entry represents the time (in seconds) it takes to run the current SINATRA algorithm based on: *(i)* the number of shapes analyzed $n = \{25, 50, 75\}$, *(ii)* the number of cones of directions $c = \{25, 50, 75\}$, *(iii)* the number of directions within each cone $d = \{5, 10\}$, and *(iv)* the number of sublevel sets (i.e., filtration steps) used to compute the Euler characteristic (EC) along a given direction $l = \{25, 50\}$. We simulate 10 different datasets for each combination of parameter values. Computations were performed with single cores on the OSCAR high performance clustering system hosted by the Center for Computation and Visualization (CCV) at Brown University. The last combination $\{c = 75, d = 10, l = 50\}$ could not be completed due to memory restrictions on the machine. Values in the parentheses are the standard deviations of these estimated times across the different runs.

| | Test | Region Size | *Tarsius* vs. *Saimiri* | *Tarsius* vs. *Mirza* | *Tarsius* vs. *Microcebus* |
|---|---|---|---|---|---|
| *P-values (P)* | KNN | 10 | $\mathbf{2.00 \times 10^{-2}}$ | $2.77 \times 10^{-1}$ | $\mathbf{2.40 \times 10^{-2}}$ |
| | | 50 | $\mathbf{1.80 \times 10^{-2}}$ | $3.20 \times 10^{-1}$ | $\mathbf{2.00 \times 10^{-2}}$ |
| | | 100 | $\mathbf{2.20 \times 10^{-2}}$ | $3.35 \times 10^{-1}$ | $\mathbf{2.00 \times 10^{-2}}$ |
| | | 150 | $\mathbf{3.59 \times 10^{-2}}$ | $3.55 \times 10^{-1}$ | $\mathbf{3.19 \times 10^{-2}}$ |
| | | 200 | $\mathbf{6.19 \times 10^{-2}}$ | $3.41 \times 10^{-1}$ | $\mathbf{2.59 \times 10^{-2}}$ |
| | Equal-Area | 10 | $\mathbf{7.98 \times 10^{-2}}$ | $2.71 \times 10^{-1}$ | $\mathbf{9.38 \times 10^{-2}}$ |
| | | 50 | $\mathbf{2.79 \times 10^{-2}}$ | $2.71 \times 10^{-1}$ | $\mathbf{3.20 \times 10^{-2}}$ |
| | | 100 | $\mathbf{2.99 \times 10^{-2}}$ | $3.61 \times 10^{-1}$ | $\mathbf{3.20 \times 10^{-2}}$ |
| | | 150 | $\mathbf{4.39 \times 10^{-2}}$ | $3.81 \times 10^{-1}$ | $\mathbf{4.19 \times 10^{-2}}$ |
| | | 200 | $\mathbf{6.19 \times 10^{-2}}$ | $3.53 \times 10^{-1}$ | $\mathbf{6.99 \times 10^{-2}}$ |
| *Bayes Factors (BF)* | KNN | 10 | **4.709** | 1.034 | **4.116** |
| | | 50 | **5.095** | 1.009 | **4.709** |
| | | 100 | **4.388** | 1.004 | **4.709** |
| | | 150 | **3.078** | 1.001 | **3.345** |
| | | 200 | **3.078** | 1.003 | **3.882** |
| | Equal-Area | 10 | **1.823** | 1.039 | **1.657** |
| | | 50 | **3.680** | 1.039 | **3.345** |
| | | 100 | **3.502** | 1.000 | **3.345** |
| | | 150 | **2.680** | — | **2.767** |
| | | 200 | **2.137** | 1.001 | **1.979** |

**Supplementary Table 3. Null region experiment to evaluate the ability of the baseline Limit Shapes algorithm with normalized vertex weights to find paraconids in *Tarsius* molars.** Here, we assess how likely it is that the Limit Shapes algorithm finds the region of interest (ROI) by chance. We generate 500 "null" regions on each *Tarsius* tooth using *(i)* a KNN algorithm and *(ii)* an equal-area approach (Supplementary Material, Section 5). Next, for each region, we sum the evidence potential or "birth times" of all its vertices. We compare how many times the aggregate scores for the ROI is less than those for the null regions. The median of these "$P$-values", and their corresponding calibrated Bayes factors (BF) when median $P < 1/e$, across all teeth are provided above for the three primate comparisons. Results with values $P$-values less than 0.1 and BFs greater than 1.598 are in bold.
